# Unconventional Fusion Mechanism at the Origin of Eukaryotic Membranes

**DOI:** 10.1101/2025.10.07.680909

**Authors:** Maria L. Mascotti, Luis S. Mayorga, Diego Masone

## Abstract

Eukaryogenesis remains one of biology’s most intriguing transitions, yet the events driving the emergence of the eukaryotic cell membrane have not been sufficiently explored. Canonical membrane fusion models, are not appropriate to explain the transition from an archaeal cell membrane to a bacterial one, via heterochiral intermediates. Here, we show that a non-canonical, lipid-mediated mechanism spontaneously generates closed bilayers of mixed bacterial and archaeal lipids. Using as a proxy a combination of enhanced-sampling and unbiased molecular dynamics simulations, we demonstrate that transient edge-mediated archaeal intermediates merge with bacterial vesicles without the formation or expansion of a fusion pore. Key indicators include reduced energy barriers and significant membrane stability post-fusion. The edge-induced route bridges the inconsistencies found in protocell models and protein-dependent pathways, proposing an unconventional explanation that describes how early eukaryotic systems achieved membrane continuity. These findings provide a plausible biophysical basis for the origin of the eukaryotic plasma membrane.

## Introduction

It is well accepted that the Eukarya domain originated from the merger between an archaeal host and a bacterial endosymbiont. Along this event, a series of major transformations took place giving rise to the first eukaryotic cells populating earth (known as Last Eukaryotic Common Ancestor, LECA), displaying a composite structure with a defined nucleus, mitochondria and a complex endomembrane system.^1^ The plasma membrane of this lineage is characterized by ester-linked fatty acids attached to a glycerol-3-phosphate (G3P) organized in a bilayer alike the one of Bacteria domain.^2^ On the contrary, the archaeal cell membrane is typically formed by ether-linked isoprenoid chains attached to a glycerol-1-phosphate (G1P) head-group.^3^ Moreover, archaea contain tail-to-tail linked lipids known as archaeols, which transform the membrane from a bilayer into a more rigid, monolayer-like structure,^4^ thereby conferring exceptional stability under extreme environmental conditions.^5^ This key difference in membrane composition –known as the ‘lipid divide’– has been extensively investigated in the context of origin of life research.^6^

Several hypotheses of the eukaryogenesis via endosymbiosis have been described,^7^ which basically propose alternatives to the order of intracellular concurrent events.^8^ However, a common feature to all of them is the assumption that the bacterial endosymbiont ‘took over’ the membrane of the archaeal host, via the ‘natural fusion’ of produced outer membrane vesicles (OMVs) with the host plasma membrane.^9^ It has been demonstrated that mixed G3P-G1P membranes are viable in modern bacteria, implying that intermediate states are plausible.^10^

Something like a ‘natural fusion’ mechanism could be interpreted as canonical membrane fusion, however among the two lineages this seems unlikely to happen. In most organisms, the two faces of a biological membrane are exposed to significantly different environments *e*.*g*., the cytoplasm and the extracellular medium. Each side is specialized for distinct functions and therefore has a unique protein and lipid composition.^11^ Consequently, when two initially separate compartments come into contact, a membrane fusion event is required to merge them. There are two main types of membrane fusion events: one that begins with the interaction of the extracellular surfaces (such as in sperm-egg or virus-cell fusion) and another one that starts with the contact of the intracellular faces (as between intracellular organelles or during exocytosis).^12^ While membrane fusion involving either the extracellular or intracellular hemi-membranes shares similar biophysical mechanisms,^13^ the proteins involved among both processes differ significantly.^14^ The bacterial-archaeal membrane fusion event does not fit into either of these mechanisms, as the bacterial outer surface must merge with the archaeal inner one. This is non-trivial, as no mechanism has been described for such interaction.

Membrane fusion necessarily involves an important topological change between the involved membranes, requiring major lipid reorganization.^15^ Such event (*e*.*g*., during neuro-transmitter release, virus entry into host cells, or intracellular trafficking) often stands on a hemifusion intermediate, like a metastable stalk or diaphragm that precedes final full fusion.^16^ Although certain lipids and proteins are known to affect the free energy and speed of the fusion process, measuring their effects is experimentally challenging. Computational methods have identified potential hemifusion pathways and their associated free-energy landscapes, however there have been relatively few studies examining the impact of lipid composition on the fusion process.^17^ Consequently, important collective mechanisms only observable under particular combinations of lipids remain unexplained.

In this work, we employ large-scale computer simulations to uncover the emergence of the eukaryotic cell membrane during the eukaryogenesis process. Using enhanced sampling simulations, we obtained thermodynamic data for hemifusion stalks between various combinations of model bacterial and archaeal membranes. Combined with long unbiased classical molecular dynamics of vesicles under confinement (resembling the initial bacterium-inside-archaeon scenario) we analyzed their mixing. Our results reveal a fundamental incompatibility between the canonical fusion-pore model and the proposed bacterial-archaeal membrane fusion mechanism. We observe a distinct pathway driven by archaeal membrane edge instabilities invading bacterial-like vesicles, suggesting an unexpected fusion mechanism in eukaryogenesis.

## Methods

### Molecular dynamics

Unbiased molecular dynamics simulations were conducted with Gromacs.^18^ Enhanced molecular dynamics simulations were conducted with Gromacs patched with Plumed.^19^ All simulations were conducted in the Martini coarse-grained space.^20^ Systems were prepared with the Charmm-gui web server^21^ and the `insane.py` (INSert membrANE) tool.^22^ All systems were minimized and equilibrated following the Martini recommendations at www.cgmartini.nl and the Charmm-gui protocol^23^ recommended for combining Gromacs with Martini.

Parameter files for NPT production runs used the integrator set to md for standard molecular dynamics (in the semi-isotropic and isotropic ensembles, depending on the system), with a time step (dt) of 0.020ps. Simulations used periodic boundary conditions in all three dimensions (xyz) and the Verlet cutoff scheme for efficient handling of long-range interactions using a buffer tolerance of 0.005 kJ/mol/ps and a neighbor list update frequency every 20 steps. Electrostatic interactions were modeled with a reaction-field method with a relative dielectric constant of 15 and a Coulomb cutoff of 1.1nm (rcoulomb), while van der Waals interactions were also truncated at 1.1nm with the Potential-shift-verlet modifier. Temperature was controlled using the V-rescale thermostat^24^ at 323°K (well above the phase transition temperature, 314°K, for DPPC^25^), and pressure was controlled using the Parrinello-Rahman barostat^26^ with a coupling time constant of 12ps and a target pressure of 1.0bar. These settings ensured efficient and stable dynamics while maintaining physical conditions appropriate for long simulations involving membrane and solvent systems.^23^

### Membrane fusion

Here, we have followed the methodology proposed by Poojari et al.^17^ in employing a reaction coordinate for umbrella sampling, along a modeled pathway of stalk formation. However, we acknowledge that a modeled reaction coordinate does not guarantee the minimal free energy path among all possible ways to go from the initial bilayer-bilayer state to the stalk. Accordingly, Smirnova et al.^27^ reported the minimum free energy path computed via a string method, which does not predefine a specific coordinate. In close agreement with the reaction coordinate method, they found that the stalk formation pathway depends strongly on the inter-membrane distance and that the minimum free energy path reveals both the barrier and a metastable stalk.

As previously investigated,^17^ varying quantities of water molecules between the membranes lead to different equilibrium inter-membrane distances, which significantly influence the free-energy landscape for membrane hemifusion. Therefore, all systems in this study were prepared, minimized and equilibrated for the same initial inter-membrane distance of ∼ 4*nm* (measured between parallel mid-planes defined by phosphate head-groups). See density profiles in Supporting Information. Equation 1 shows the collective variable (*ξ*_*f*_) used to bias molecular dynamics simulations.

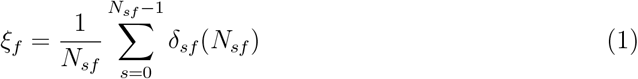

Here, *N*_*sf*_ is the number of tail beads inside slice *s* of the cylinder. *δ*_*sf*_ is a continuous function of *N*_*sf*_ within the interval [0 1] taking the values: *δ*_*sf*_ =0 for no beads in slice *s* and *δ*_*sf*_ =1 for at least one bead in slice *s*. For mathematical details see supplementary information and the original article.^28^ The fusion process begins with a pair of flat and parallel independent bilayers (0.2 *< ξ*_*f*_ *<* 0.3). The first hemifusion stalk forms at 0.5 *< ξ*_*f*_ *<* 0.7 and then it grows until *ξ*_*f*_ ∼ 1. The collective variable pulls from tail beads (C4A & C4B for DPPC and C4C & C4D for bolalipids) to fill a cylinder with *N*_*sf*_ slices, of thickness *d*_*sf*_, radius *R*_*cylf*_ and an occupation factor *ζ*_*f*_ . Importantly, *ξ*_*f*_ enforces connectivity between the two hydrophobic cores without constraining a priori the detailed shape or composition of the stalk.

Theoretically, the free energy required to transition between different thermodynamic states (*i*.*e*., parallel bilayers to first hemifusion) should be independent of the direction of the collective variable.^29^ Consequently, the formation pathways of the hemifusion stalk should exhibit identical free-energy profiles, regardless of the direction: whether inducing a stalk from separated bilayers or unfusing bilayers from an already formed stalk. Any substantial differences indicate potential issues with hysteresis, insufficient sampling, or poor convergence. Collective variable *ξ*_*f*_ has shown negligible hysteresis effects during membrane fusion,^17^ which we verified by conducting forward and backward biased simulations^30,31^ as originally suggested for free-energy calculations.^32^ Moreover, *ξ*_*f*_ was extensively validated, demonstrating its applicability among lipid mixtures,^17^ combined with nanoparticles,^33^ assessing finite-size effects,^17,34^ and benchmarking it against earlier reaction coordinates.^28,35^

### Potential of mean force

Free-energy profiles were calculated using umbrella sampling^36^ in Plumed^19^ and analyzed with the Weighted Histogram Analysis Method (WHAM).^37^ To cover the hemifusion stalk free-energy profiles, 17 windows were used to sample collective variable *ξ*_*f*_ within the range of ∼[0.2, 1]. A force constant of *k* = 50, 000*kJ/mol* was applied in all cases. Each window used for recovering the free-energy profile included at least 100 ns in the steady-state regime, any transient data were removed from the analyses. More technical details on umbrella sampling for the stalk formation using PLUMED are provided in our previous works.^30,31^ The membrane fusion collective variable is compilable from PLUMED-v2.9 by adding `–enable-modules=membranefusion` to the `configure` command. For more details see: www.plumed.org/doc-v2.9/user-doc/html/_m_e_m_b_r_a_n_e_f_u_s_i_o_n_m_o_d.html

### Vesicle preparation

To prepare Gromacs-ready-to-simulate vesicle-inside-vesicle systems,^38^ we first generated two symmetric bilayers with the `insane.py` (INSert membrANE) tool^22^ and disposed them at the desired inter-membrane distance (∼ 4*nm*) to produce a double-bilayer structure. We used BUMPy (Building Unique Membranes in Python)^39^ to generate a spherical membrane geometry and to create a vesicle-within-vesicle arrangement. BUMPy builds spherical vesicles by assigning lipids to each monolayer based on the monolayer pivotal plane. Starting from a pair of flat bilayers, a vesicle midpoint radius is then chosen to estimate the pivotal plane depth. BUMPy then computes the pivotal-plane outer and inner areas and sets the number of lipids accordingly, so each leaflet in the vesicle-in-vesicle arrangement starts with approximately the same intrinsic area-per-lipid as the flat reference. We solvated the composite structure using `insane.py` and standard Martini W water molecules. Although we initially built several systems of different sizes, we systematically observed effective active-edge fusion and hemifusion diaphragm formation for ∼1,600 DPPC vesicles confined in ∼1,800 BOLB ones, all solvated in ∼ 260, 000 water molecules in a ∼30x30x30nm cubic box.

As we observed before,^40^ when two lipid species have markedly different tail lengths, tight membrane curvature can produce pronounced local asymmetries: shorter-tailed lipids preferentially concentrate in highly curved regions, while longer-tailed lipids migrate toward areas of lower curvature. This curvature-driven lipid sorting has been described before.^41–43^ As we discussed in a previous study on DOPC/DTPC vesicles under confinement,^40^ shape-induced lipid relocation relies on the understanding that curvature is a collective property,^44^ meaning that the orientational stress experienced by each lipid molecule depends not only on its own orientation but also on the orientations of its neighbors. There, we reported curvature values up to 0.16*nm*^−1^ for highly confined inner micelles. Here, we built vesicles that could in principle realistically mimic the experimental curvature: reported values for OMVs range in diameter from 10 to 450nm (90 to 450nm,^45^ 50 to 300nm,^46^ and 10 to 300nm^47^). However, to minimize computational costs still within the reported values, we chose outer vesicles of ∼30nm in diameter containing half size inner ones. Such geometry yields ∼53,000 W beads in the confined space between vesicles.

Following the same procedure for vesicle preparation and systems minimization and equilibration, we also simulated phosphatidylethanolamine (DPPE) and dioleoylphosphatidylglycerol (DOPG) for inner vesicles, keeping bolalipids in the outer container. For BOLB:DPPE systems, we used ∼1,800 BOLB and ∼1,900 DPPE solvated in ∼ 260, 000 water molecules in a cubic box of ∼33x33x33nm. For BOLB:DOPG systems, we used ∼1,800 BOLB and ∼1,950 DOPG solvated in ∼ 260, 000 water molecules with sodium ions (*Na*^+^) to neutralize the total charge, also in a cubic box of ∼33x33x33nm.

### Post-processing and graphics

Leaflet segmentation was conducted with MDVoxelSegmentation^48^ and object completion was applied with MDVWhole.^49^ MDVContainment^50^ was used to identify hierarchies of confined water cavities.

Figures were made with Visual Molecular Dynamics (VMD),^51^ Grace (GRaphing, Advanced Computation and Exploration of data),^52^ Inkscape^53^ and Octave.^54^ Videos were rendered with Ovito.^55^

## Results

### Bolalipids increase the energy cost for membrane hemifusion

As a proxy to construct a system resembling archaeal membranes, we employed double-headed bolalipids. In the Martini force-field,^20^ bolalipids consist of two dipalmitoylphosphatidylcholine (DPPC) lipids covalently linked together at either one (acyclic, labeled BOLB) or both tail ends (cyclic, labeled BOLA).^56^ Therefore, we prepared bilayers composed of DPPC, BOLA and BOLB in different proportions to study membrane hemifusion. Table 1 lists all systems considered.

**Table 1:** Pairs of bilayers for hemifusion.

| system # | *upper membrane* | *lower membrane* |
| --- | --- | --- |
| 1 | DPPC | DPPC |
| 2 | DPPC:BOLA (90:10) | DPPC:BOLA (90:10) |
| 3 | DPPC | BOLA |
| 4 | BOLA | BOLA |
| 5 | DPPC:BOLB (90:10) | DPPC:BOLB (90:10) |
| 6 | DPPC | BOLB |
| 7 | BOLB | BOLB |

To effectively induce the formation of the hemifusion stalk, we followed a previously developed methodology^30^ that biases molecular dynamics simulations using the parameter *ξ*_*f*_, which quantifies the degree of fusion stalk formation (see the Methods section for full technical details on membrane fusion using enhanced sampling).

Fig. 1a shows the free-energy profiles to induce a hemifusion stalk using umbrella sampling.^36^ All systems started from initially flat and parallel membranes at 0.2 *< ξ*_*f*_ *<* 0.3. The first stalks forms at 0.5 *< ξ*_*f*_ *<* 0.7, while for *ξ*_*f*_ *>* 0.7 they widen. Pure bolalipid membranes (cyclic and acyclic) exhibit the highest energy cost for the stalk formation and its further enlargement. On the contrary, pure DPPC bilayers show the lowest energy costs. Intermediate cases include combinations of DPPC and bolalipids, namely: one DPPC bilayer and one bolalipid bilayer and bilayers containing 90% of DPPC and 10% bolalipids.

**Figure 1:**
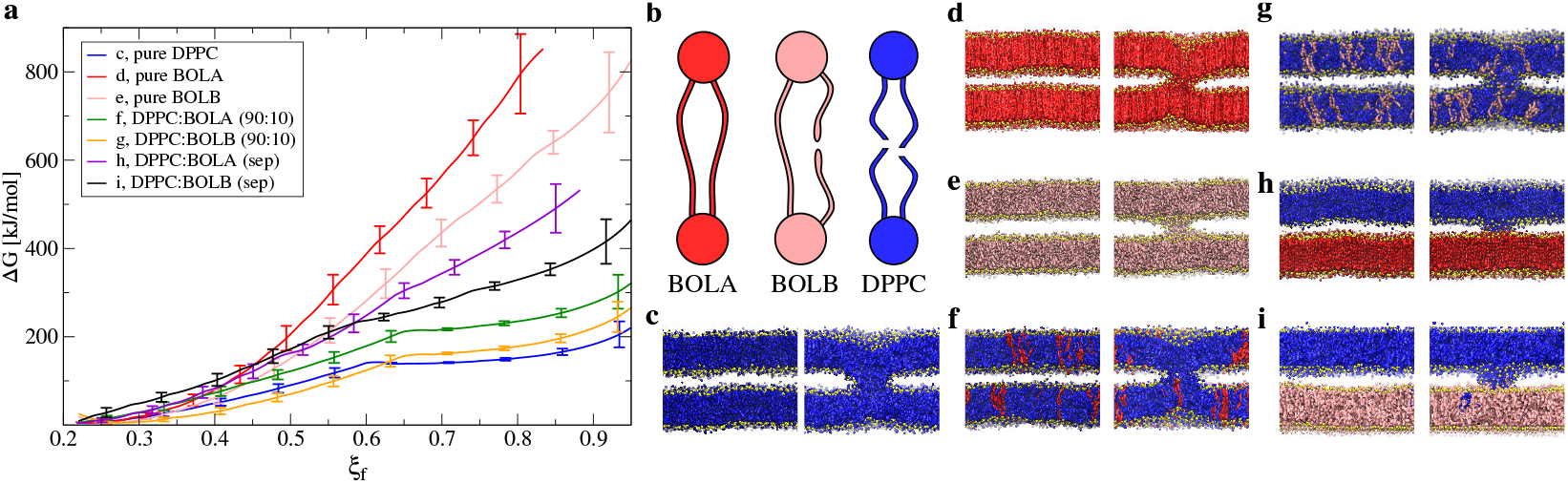
Thermodynamics of hemifusion stalk among bacteria and archaea. **a**, Potential of mean force. Error bars are standard errors calculated by individually splitting the profiles in independent blocks. **b**, Drawings of cyclic bolalipids (BOLA) in red, acyclic bolalipids (BOLB) in pink and standard dipalmitoylphosphatidylcholines (DPPC) in blue. **c-i**, Molecular dynamics snapshots show the initial (left) and final (right) states of process during stalk formation along parameter *ξ*_*f*_ . DPPC is blue, BOLA is red, BOLB is pink and phosphate groups (PO4, PO1 and PO2) are yellow. For clarity, water molecules are not shown.

Although stalks successfully form in all cases, significant structural differences are observed. First, pure and identical membranes (Fig. 1c-e) show that DPPC bilayers form a wider stalk than bolalipid ones, which are slightly tilted. Second, for systems containing 90% of DPPC and 10% of bolalipids (Fig. 1f,g) the stalk is mainly composed of DPPC. Third, Fig. 1h,i shows that when confronted with a pure bolalipid membrane, the DPPC bilayer takes over the stalk. Moreover, Fig. 1h shows how some DPPC lipids have even diffused into the bolalipid membrane (for intermediate states snapshots please see Fig. S1).

In simulations, the short-range hydration repulsion between two apposed membranes (or between a membrane and its periodic image) may contribute with a free-energy penalty that manifests as an additional surface tension.^57^ Therefore, we quantified whether bolalipids are likely to enhance hydration repulsion, thus providing an additional mechanistic argument against traditional stalk formation. Following the methodology described by Smirnova et al.^57^ we estimated the hydration repulsion in *logP* ∼ 5mN/m, in agreement with reported values for other model membranes.^57,58^ See Supporting Information for calculation details.

Also, prior computational works showed that Δ*G*_*stalk*_ rises very steeply with the intermembrane distance.^57,59^ Moreover, under high-hydration conditions the stalk ceases to be metastable showing no local minimum, whereas under low-hydration conditions a metastable or even a global-minimum stalk may appear.^17^ In addition, lipid composition and the choice of coarse-grained force-field substantially modulate absolute free energies. Therefore, free-energy values for stalk formation should be read here as a condition-specific result that illustrates trends rather than universal biological constants.^17^

With all systems initially fixed to the same inter-membrane distance (∼ 4*nm*), free energies for stalk formation in Fig. 1 suggest that archaea:bacteria hemifusion seems significantly less probable than bacteria:bacteria hemifusion. Moreover, long unbiased molecular dynamics simulations of parallel membranes reveal that the hemifusion stalk is a low-probability event, meaning that except for very low hydration states, hundreds of microseconds can be simulated without observing any hemifusion at all. This seems to be true at least for the accepted mechanism of membrane fusion, which starts with the formation of the hemifusion stalk, follows with the nucleation and radial expansion of a fusion pore, to finally achieve full-fusion when the content of the up and down compartments mix.^60^ Consequently, hemifusion between archaea and bacteria seems to be unlikely, suggesting that another mechanism to mix their membranes must exist.

### Vesicles in confinement reveal bacterial-archaeal mixing

We have previously observed that spatial confinement —such as that created when an outer vesicle engulfs an internal compartment— can trigger unexpected membrane interactions.^40^ From the perspective of the endosymbiotic origin of eukaryotes,^61^ this is particularly relevant: eukaryotic cells are thought to have arisen when an archaeal host engulfed an aerobic bacterium, which ultimately evolved into mitochondria and contributed to the emergence of the endomembrane system via the production of OMVs that fused with the host’s plasma membrane. The three-dimensional dynamics of such a scenario suggest that spatial confinement could play a critical role. However, how confinement influences the interaction between OMVs and archaeal membranes remains unexplored.

To address this first problem, we constructed several model bacterial vesicles, confined by archaeal ones^56^ using different proportions of bolalipids and dipalmitoylphosphatidylcholines, as listed in Table 2. We conducted dozens of simulations in the *µ*s length-scale at different sizes, to account for diverse degrees of initial confinement. As expected, large outer vesicles relative to the inner ones (*>*2:1 diameter ratio), imposed minimal spatial restrictions, making fusion events unlikely. Therefore, such systems showed no lipid mixing between the container and contained vesicles, regardless of the lipid composition.

**Table 2:** Pairs of vesicles for hemifusion.

| system # | *inner vesicle* | *outer vesicle* | *mixing* |
| --- | --- | --- | --- |
| 1 | DPPC | BOLA | no |
| 2 | DPPC | DPPC:BOLA (90:10) | no |
| 3 | DPPC | DPPC:BOLB (90:10) | yes |
| 4 | DPPC | DPPC:BOLB (50:50) | no |
| 5 | DPPC | BOLB | yes |

In principle, by increasing confinement (∼1.25:1 diameter ratio), lipid mixing is expected to become more probable. However, only systems #3 and #5 showed lipid mixing, both containing acyclic bolalipids. Having just 10% of bolalipids, system #3 derived into a stomatocyte-like structure identical to the ones previously observed between confined phosphatidylcholines vesicles.^38,40^ On the contrary, system #5 evolved into an unexpected hemifusion intermediate in the shape of a diaphragm, consistently over multiple trajectory repetitions (n=10, as recommended for molecular dynamics to avoid false positive conclusions^62^). The other systems (#1,#2 and #4) showed structures that systematically avoided fusion events (see Figs. S2-S4).

### Active edges enable bacterial-archaeal hemifusion

A careful analysis of the trajectory repetitions in system #5 unveils a common mechanism for the formation of the hemifusion configuration. Fig. 2 shows a DPPC vesicle (blue) confined in a BOLB container (pink) as they evolve in time during one of the 10*µ*s unbiased simulations (see Fig. S5 for more snapshots). First, the BOLB vesicle opens multiple membrane pores which expand and mix between themselves, while the inner DPPC vesicle takes an oblate shape (Fig. 2b,c). Once the container reorganizes into a cup-shaped structure (Fig. 2d), its edges remain highly unstable as bolalipids are difficult to accommodate in such high-curvature regions, here termed active edges. Over multiple repetitions of this trajectory (n=10), active edges systematically fuse to the inner DPPC vesicle (Fig. 2e), to form the intermediate hemifusion structure, in the shape of a diaphragm (Fig. 2f). At this stage some bolalipids have diffused into the external hemilayer of the DPPC vesicle and the two water cavities are separated by a single DPPC bilayer, in agreement with a diaphragm configuration.^63^

**Figure 2:**
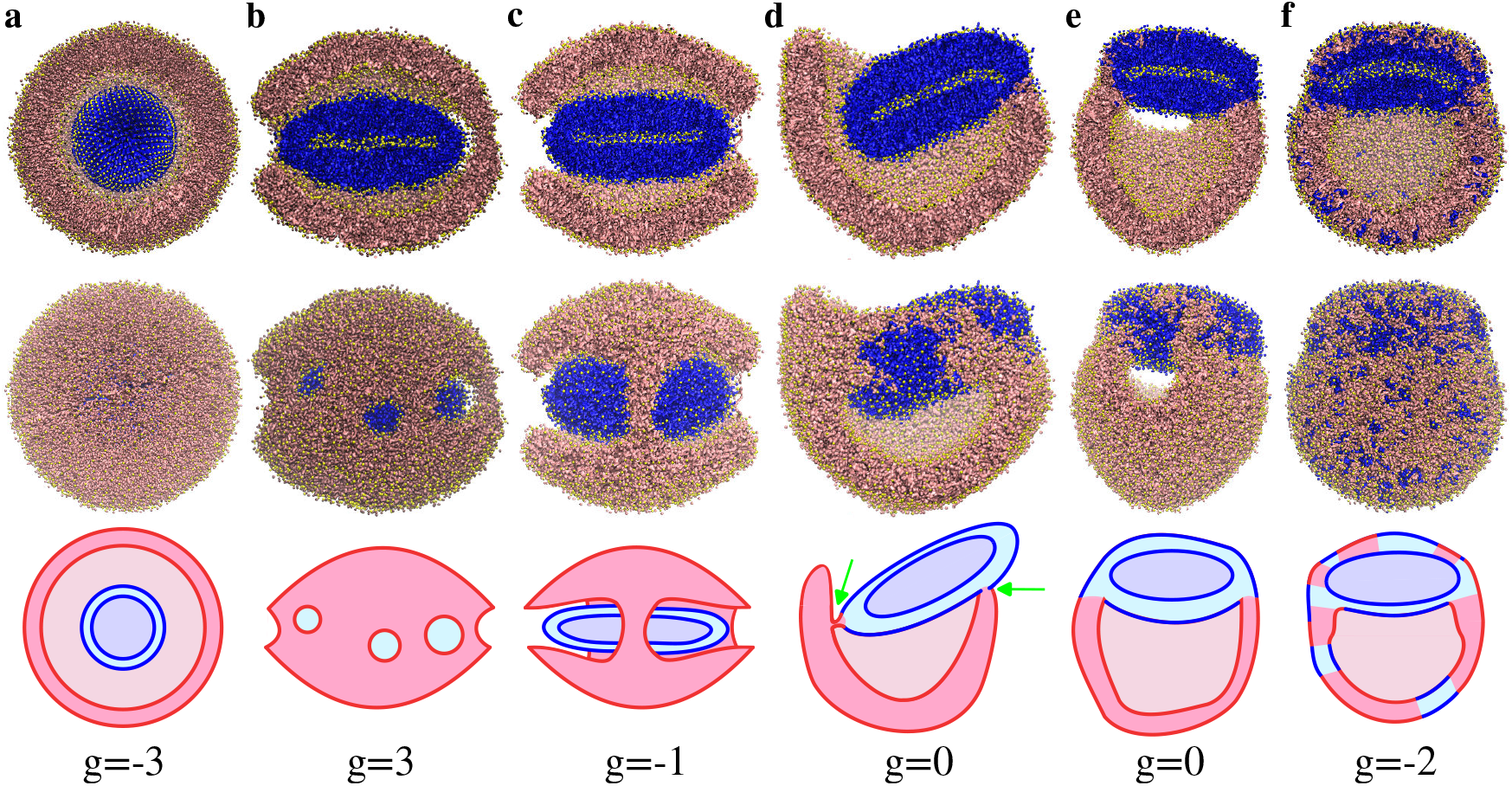
Bolalipid active edges dock with the dipalmitoylphosphatidylcholine vesicle. **a-f**, Snapshots depicting the full fusion process over time, proceeding via an acyclic bolalipid active edge mechanism. **Top row:** Cross-section. **Middle row:** Outside view. DPPC lipids are blue, BOLB are pink with all phosphate groups are yellow. **Bottom row:** Schematic drawings for genus calculation. Solid lines represent phosphate groups surfaces, in pink for bolalipids, and in blue for dipalmitoylphosphatidylcholines. Green arrows indicate selected topological events.

To follow the complexity of the 3D structures derived, we use the genus as a topologically invariant property along molecular dynamics trajectories to identify changes over time. As suggested for lipid vesicles,^64^ the genus *g* is defined by the equation:

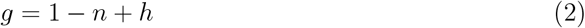

where *n* is the number of non-intersecting surfaces (inner/outer faces) and *h* is the number of holes (pores). From the initial structure with four independent surfaces (internal and external faces of the two concentric vesicles, *g* = −3, Fig. 2a) the outer vesicle forms several unstable pores that transiently increase the genus (*g* = 3, Fig. 2b). The process converges into the basket-like structure with *g* = 0 (the inner surface of the DPPC vesicle and the outer surface covering all the external face of the structure, with a large pore under the handle, Fig. 2d). The closing of the basket renders the hemifusion diaphragm by sealing the active edges at both sides of the handle, with three independent surfaces and no pores: one external facing the bulk water and two internals surrounding the two confined aqueous compartments (*g* = −2, Fig. 2f).

Experimental observations indicate that unstable edges strongly promote interaction and fusion with nearby liposomes.^65^ Also, large hemifusion intermediates have been reported when influenza virus-like particles interact with liposomes.^66^ The authors propose that viral hemagglutinin drives membrane merging via two independent pathways: a rupture-insertion mechanism that produces extended hemifusion diaphragms —analogous to those described here— and the classical stalk-hemifusion-fusion-pore route. In the 2D schematics of the rupture-insertion pathway, formation of a large hemifusion diaphragm appears straightforward: the U-shaped edges of the virus-like particle attach to the liposome membrane, creating two closed compartments separated by a bilayer. However, in 3D this process demands extensive lipid reorganization as previously reported in computational studies of hemifusion diaphragms stabilization.^67,68^ Here, in the initial vesicle-in-vesicle arrangement, membrane rupture yields a cup-shaped structure as observed for bolalipid membranes (Fig. 2c,d). Transitioning to a hemifusion diaphragm therefore requires a substantial topological transformation, characterized by a change in genus from g=0 to g=-2 (Fig. 2e,f). This remodeling involves the closure of two membrane pores, sealing the outer surface, restoring membrane continuity, and producing two closed compartments separated by a contiguous bilayer.

### The active-edge fusion pathway

Using self-consistent field theory it has been shown that an elongated worm-like stalk adjacent to a membrane pore can reduce the effective line tension of the edge portion in contact with the stalk, relative to a bare pore, thereby lowering the activation barrier for nucleation of the stalk-pore complex.^69^ Also, previous Monte Carlo simulations unveiled a non-standard expansion of the stalk encircling a membrane pore, which drives membrane fusion and allows for transient leakage.^70,71^ These findings support active-edge fusion between vesicles: after the first fusion (Fig. 2d,e), the stalk zip-closes (elongates) effectively reducing the genus from g=0 to g=-2 (Fig. 2e,f).

The latter zip-closure of the stalk implies a transient leaky fusion mechanism, where water cavities are not always isolated during the process. Whether transient leakage is an intrinsic feature of membrane fusion remains unresolved,^70,72^ with experimental data reporting fusion with^73^ and without^74^ leakage among biological membrane fusion processes. The active-edge induced fusion mechanism proposed here, is a pathway of membrane fusion in which the rim (or edge) of an open vesicle is destabilized (Fig. 2d), driving the transition into a hemifusion diaphragm (Fig. 2e-f). Unlike a smooth and well-sealed pore formation, this edge-driven process is inherently leaky, as water content from the container vesicle escapes toward the exterior (Fig. 2b-e), before closing definitively (Fig. 2f). However, we have observed that the content of the inner vesicle remains isolated until the rim pore opens at the hemifusion diaphragm (see Fig. S6).

We also verified the active-edge fusion mechanism among systems with a pure DPPE vesicle confined by a BOLB container, which similarly to DPPC:BOLB ones, evolved into structures with a hemifusion diaphragm (see Fig. S7). On the contrary, pure DOPG vesicles showed no hemifusion intermediates (see Fig. S8), analogously to other vesicle-in-vesicle arrangements from our previous studies.^38,40^

### The lateral expansion of a rim pore marks the beginning of full fusion

To achieve full fusion, the diaphragm must disappear. Unbiased molecular dynamics naturally resolved the metastable hemifusion diaphragm observed in Fig. 2f. Graphically, Fig. 3a-d details subsequent events when a membrane pore opens near the rim of the hemifusion diaphragm,^75^ connecting the two water cavities. Counterintuitively, the membrane rim pore does not grow radially into a fusion pore,^76,77^ instead it propagates laterally along the interface border. Such perimeter-driven expansion continues until both pore ends meet for a fission event. The final result is that an inner DPPC micelle separates from the rest of the structure. Importantly, of the ten trajectories that evolved into a hemifusion diaphragm (on the tens of *µ*s time scale, see Fig. S5 and supplementary video 1), nine exhibited rim-pore opening, and eight underwent lateral expansion ending in membrane fission and release of the inner micelle into the container vesicle (see supplementary video 2).

**Figure 3:**
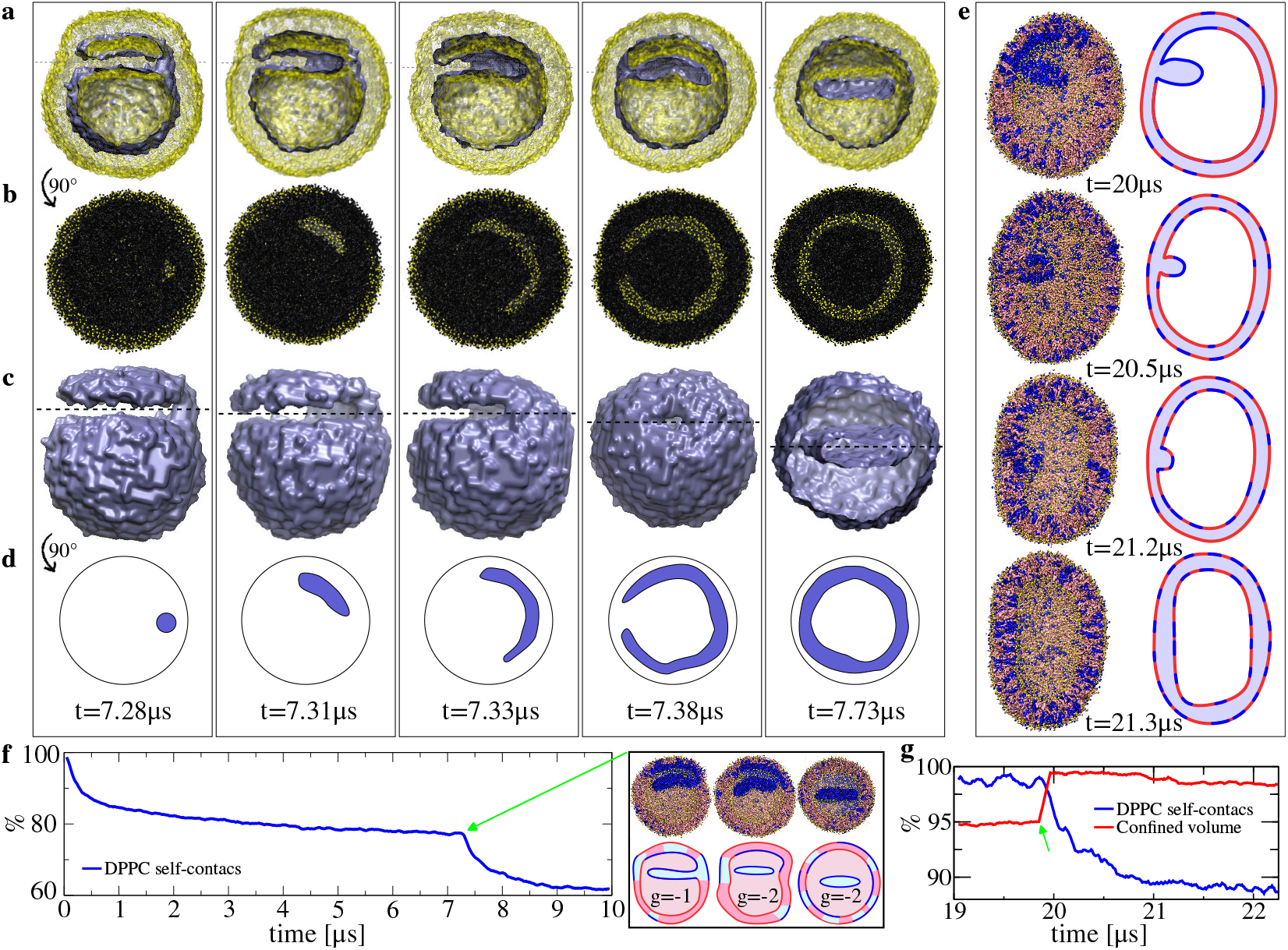
Disassembly of the hemifusion diaphragm in the progression to full fusion. **a**, Snapshots from the initial membrane rim pore through its widening along the border, up to the final scission. Water surface is shown in light purple, with phosphate beads in yellow. Dashed lines indicate planes of section for snapshots shown in row b. **b**, Cross-section with lipid beads in black except for phosphate beads which are yellow. Water is not shown. **c**, Water surface. Lipids are not shown. Dashed lines indicate planes of section for drawings given in row d. **d**, Schematic drawings of the membrane pore expansion along the rim, showing only its water channel. Time stamps indicate the simulation times. **e**, A bleb-shaped deformation fused to the inner hemilayer is slowly reabsorbed by the vesicle, showing molecular dynamics snapshots (left) and drawings (right). DPPC lipids are blue, BOLB are pink and all phosphate groups are yellow. Time stamps indicate total simulation times. **f**, Normalized DPPC self-contacts (<0.6nm) for events in panels a-d. Inset highlights the lateral expansion of the rim pore and its genus. **g**, Normalized DPPC self-contacts (<0.6nm) and normalized confined volume for events in panel e. Green arrows indicate the mixing of the two cavities.

From simulation time stamps in Fig. 3a-d it can be observed that the rim pore expansion along the perimeter of the diaphragm is relatively fast (∼ 0.2*µ*s) compared to the fission event happening ∼ 0.35*µ*s later. Fig. 3f plots normalized DPPC self-contacts as a function of time. At the beginning (t=0*µ*s) there is an inner DPPC vesicle inside a bolalipid container, marking the maximum of DPPC:DPPC contacts. As the system evolves, DPPC lipids slowly diffuse into the bolalipid container, with drastic changes of slope indicating major topological/morphological events. Inset in Fig. 3f highlights the change of genus from the opening of the rim pore, its lateral expansion and the fission event that excises the inner micelle. Therefore, full fusion takes place here via an unconventional mechanism, with both vesicles merging seamlessly into a continuous membrane, while leaving behind a pure DPPC micelle. See Tables S1 and S2 for statistics on these events along trajectory repetitions.

### The puzzle of diaphragm assembly and disassembly

The proposed mechanism involves the formation of relatively large diaphragm-structures that have been experimentally observed during protein-free fusion of giant unilamellar vesicles^78^ and yeast vacuoles.^79^ However, the occurrence of such expansive structures entails mechanistic processes that remain poorly understood. In the canonical stalk-to-diaphragm transition, the expansion of the diaphragm would require either an increase in lipid content in the inner hemilayers or a corresponding reduction in the outer ones, to accommodate the growing surface area of the bilayer that separates the two compartments.^78,79^ Several hypotheses have been proposed to explain the formation of large diaphragms, particularly in lysosomal systems that include the involvement of lipid transporters or lipid delivery from the lumen.^79^

Remarkably, in our vesicle-in-vesicle simulation model we constantly observed the active edge-driven assembly of metastable hemifusion diaphragms between archaeal-like and bacterial membranes. This mechanism avoids the inconsistencies associated with asymmetries in lipid distribution at the vertex zone. The ability of active edges to dock with the bacterial vesicle, followed by the closure of the surrounding container vesicle, results in a diaphragm configuration solely dependent on the initial size of the vesicles.

The disassembly of the diaphragm by radial expansion of a pore is problematic as well. In that course of action, lipids must pack onto the inner face of the vesicle, generating a large asymmetry between its inner and outer leaflets. In our systems we identify two paths for the resolution of diaphragms, one fast (Fig. 3a-d) and one slow (Fig. 3e), comparatively. Along the slow route, taking more than 1.3*µ*s for complete reabsorption, a vesicle with a bleb-like micelle protrusion fused to the inner hemilayer is observed at t=20*µ*s (Fig. 3e). Such configuration is a byproduct of an incomplete laterally-expanded rim pore, lacking the final fission event. The delay in resolving such a system under confinement is clear: the vesicle cannot incorporate an arbitrary number of lipids into its inner hemilayer. Although the surface-area to volume relation imposes limitations,^80^ still the vesicle may change of shape (like the spheroid-to-oblate,^40^ Fig. 3e) to accommodate more/less molecules at its inner/outer leaflets.^81^ Alternatively, lipid flip-flop could solve the problem by changing the stress asymmetry between the two hemilayers.^82^ However, both solutions require significant energy barriers to be overcome, thereby making the reabsorption of the micelle a metastable process with a relatively slow transition time.^83^

On the contrary, along the fast route (Fig. 3a-d) taking less than 0.5*µ*s to completion, the entire process does not require large amounts of lipid flip-flop (the inner micelle remains pure DPPC, inset in Fig. 3f), nor major shape changes to compensate for surface asymmetries (the container vesicle remains spheroidal). The main energetic costs are associated to three distinctive events (Fig. 3a-d): (i) the opening of the rim pore (t= 7.28µs), (ii) its lateral expansion (7.31*µ*s < t < 7.38*µ*s), and (iii) the fission event that excises the inner micelle (t=7.73*µ*s). Since the diaphragm is metastable, its disassembly begins with the formation of a rim pore at its edge, in agreement with previous theoretical^63^ and computational studies.^75^ Full fusion may proceed via radial pore expansion, but this would require the reabsorption of the entire diaphragm, an energetically feasible event only for small diaphragms or infinite bilayers. Under confinement, the most plausible solution is the lateral expansion of the rim pore ending in a fission event that completes full fusion.

### Cholesterol negatively regulates active edge function

Although sterols are present in some bacteria,^84^ they are not as abundant as in eukaryotic membranes. Cholesterol has been observed to modulate the propensity to form large diaphragms during fusion events among influenza virus and liposomes. At high concentrations (40%) it blocks the edge-mediated mechanism for diaphragms formation promoting the classical expansion of fusion pores.^66^ Therefore, to study the active edge fusion mechanism in the presence of cholesterol, we constructed systems containing a micelle composed entirely of acyclic bolalipids, confined by an outer vesicle containing different proportions of DPPC and cholesterol (100:0, 80:20, 70:30, and 60:40) under various degrees of confinement (∼1.25:1, ∼1.5:1 and ∼2:1 diameter ratios, see Fig. S9). Fig. 4a shows a pure DPPC vesicle (blue) containing an acyclic bolalipid micelle (pink) while its highly unstable edge subsequently fuses at different locations with the inner hemilayer. Analogously, Fig. 4b shows the same acyclic bolalipid micelle now contained by a DPPC:CHOL (70:30) vesicle. Visual inspection already suggests that for 30% of cholesterol active-edge-induced membrane fusion is less effective.

**Figure 4:**
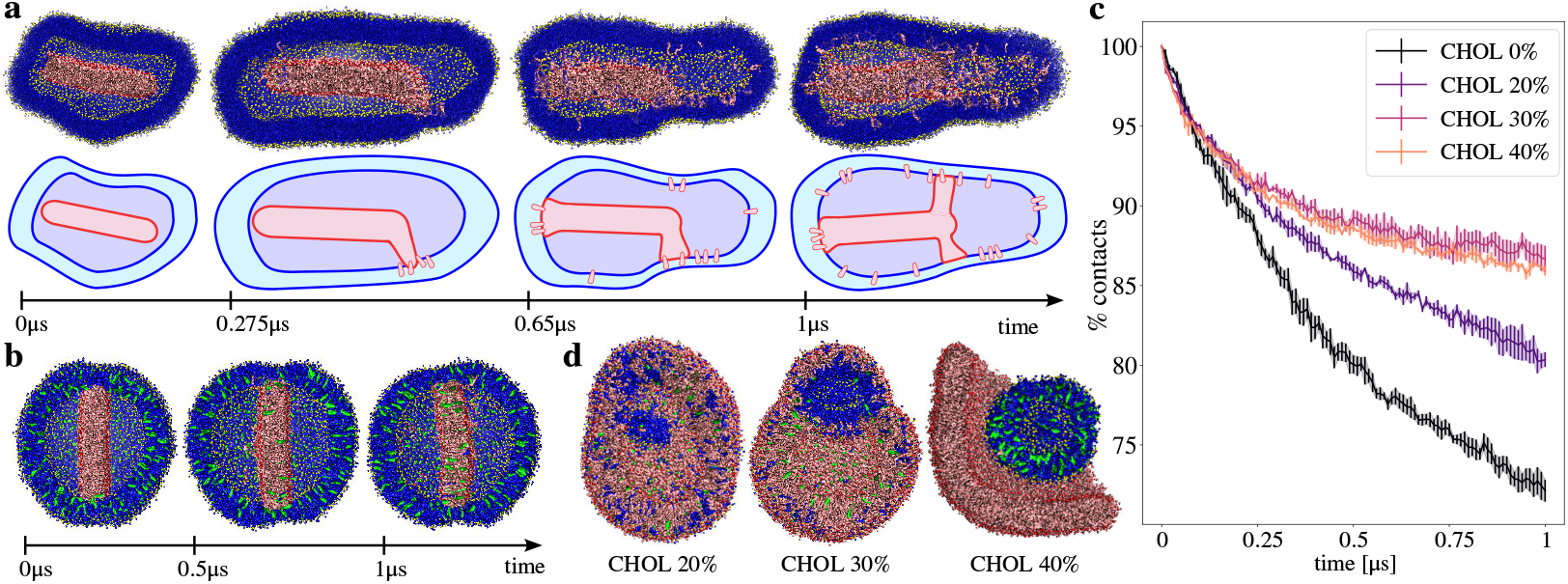
Lipid mixing is negatively regulated by cholesterol. **a**, DPPC vesicle confining an acyclic bolalipid micelle. Top: molecular dynamics snapshots and bottom, schematic drawings of the process. DPPC is blue with phosphate groups in yellow. BOLB are pink with phosphate groups in red. Water molecules are not shown. Solid lines represent phosphate groups surfaces, in pink for bolalipids, and in blue for DPPC. **b**, DPPC + cholesterol (70:30) vesicle confining an acyclic bolalipid micelle. Cholesterol molecules are green. **c**, Normalized bolalipid self-contacts at the inner micelle, averaged over trajectories repetitions (n=3) and for different proportions of DPPC + cholesterol in the outer vesicle (100:0, 80:20, 70:30 and 60:40). **d**, Molecular dynamics snapshots of vesicle-in-vesicle configurations (DPPC+CHOL inside BOLB) after 10*µ*s of unbiased simulations, for different amounts of cholesterol in the inner vesicle.

Although in all cases the inner micelle eventually docks with the outer vesicle, we observed that cholesterol reduces the ability of bolalipids to diffuse into the outer vesicle, hence attenuating active edges fusion activity. As a measure of such phenomenon, we calculated the amount of bolalipid self-interactions to quantify the integrity of the micelle over time (Fig. 4c). In absence of cholesterol, bolalipid self-contacts show the fastest decay, meaning that the micelle effectively docks with the vesicle and bolalipids diffuse into it, while increasing amounts of cholesterol progressively reduce this phenomenon (until saturation at range 30%-40%).

The effect of cholesterol reported here reflects changes in the rates of lipid mixing rather than changes in the thermodynamic driving force of membrane fusion. Namely, increasing cholesterol concentration in the vesicles slows down the observed lipid-mixing kinetics. Cholesterol is well known to increase membrane order^85^ and effective viscosity,^86^ making the concentration-dependent slowing effect a possible consequence of reduced lipid mobility, in agreement with experimental observations.^66^

Finally, we simulated confined bacterium-inside-archaeon systems (analogously to the previous sections) but now containing 20%, 30% and 40% of cholesterol in the inner vesicle. We observed that bolalipid active edges still induce lipid mixing but only at the lowest proportions of cholesterol, 20% and 30%, with no membrane fusion events observed at 40% (Fig. 4d). Also, Fig. S10 shows temporal series of events, confirming that cholesterol strongly delays the formation of the hemifusion diaphragm. Therefore, in agreement with the experimental data,^66,87^ we propose that cholesterol inhibits lipid mixing by negatively regulating active-edge-induced membrane fusion. These results suggest that during eukaryogenesis, the lipid composition of the membrane from the bacterial endosymbiont —likely containing less than 40% sterols^88,89^— may have been prone to the active-edge fusion mechanism.

## Discussion

The emergence of the eukaryotic lineage was the result of the complex merge of cell components from the two parental lineages, a bacterium and an archaeon, followed by successive stabilization/selection steps.^8^ The hypothesized fusion event among the outer surface of bacterial vesicles (likely OMVs) with the archaeon inner face of the plasma membrane cannot be explained by the canonical fusion mechanisms. Cell-to-cell or organelle-organelle fusion events share analogous structural and functional characteristics, as both are driven by anchored single-transmembrane domains, namely SNARE proteins for intracellular membrane mixing (also for exocytosis) or viral fusogens in cell infection by enveloped viruses. During conventional exocytosis, the cytoplasmic face of a vesicle contacts the cytoplasmic face of the plasma membrane. Upon fusion, the vesicle membrane is incorporated into the plasma membrane and the lumen becomes part of the extracellular space. In such a process, all the membrane components including associated and transmembrane proteins, preserve their original orientation. However, the proposed fusion of OMVs to the plasma membrane of the archaeal host presents a serious inconsistency. In a simplified configuration with a bacteria confined by an archaeon (Fig. 5a), upon fusion any OMV membrane associated component, such as ion channels or single-transmembrane proteins, would end up facing the opposite side of the resulting cell. This would generate an inversion of the membrane configuration, making it unlikely (Fig 5a).

**Figure 5:**
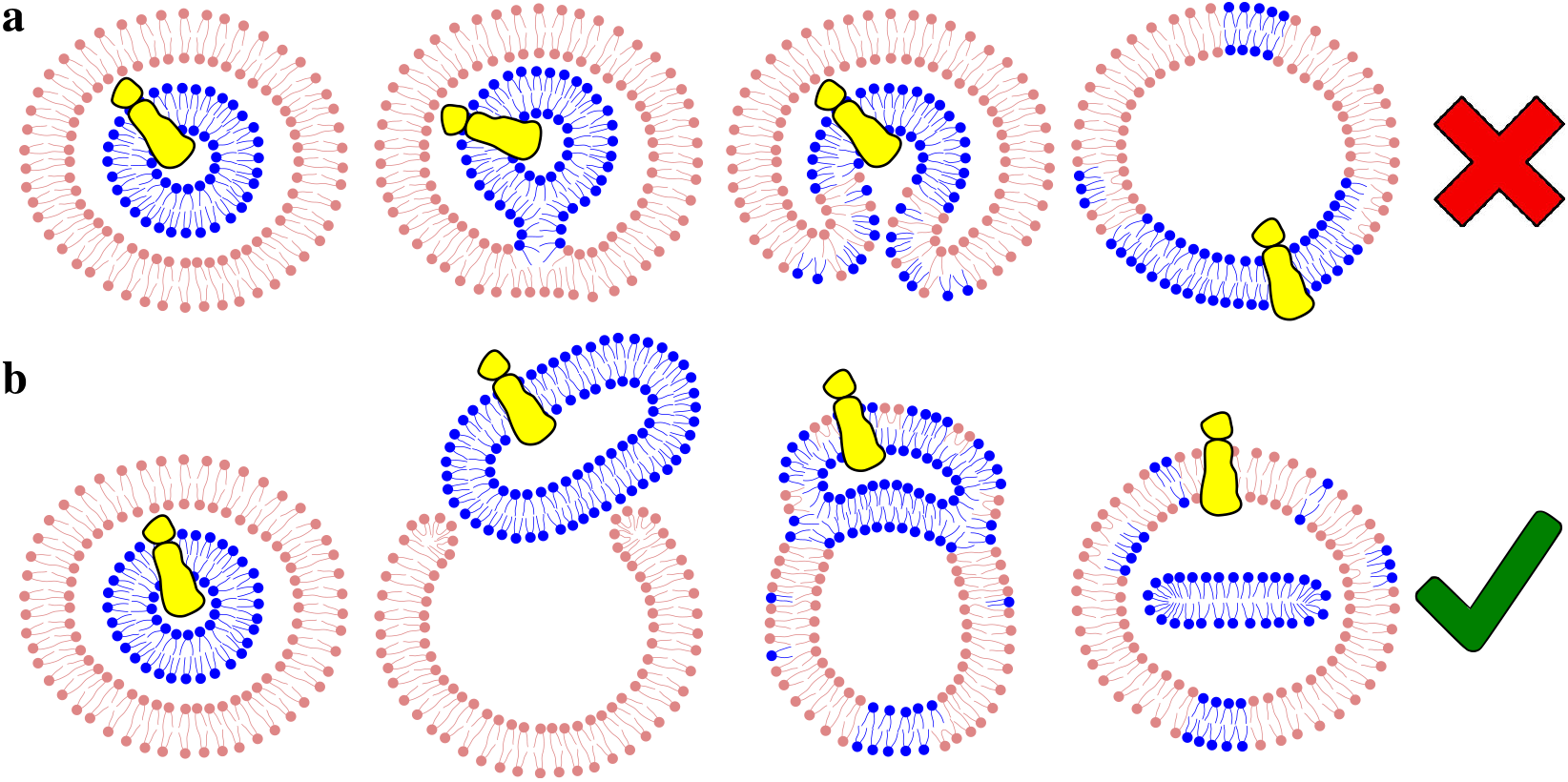
Schematic drawings of the possible archaeal-bacterial fusion mechanisms during eukaryogenesis. **a**, Canonical membrane fusion via a hemifusion stalk and a fusion pore. DPPC lipids from bacteria are colored in blue while bolalipids from archaea are pink. Yellow structures indicate a hypothetical directional transmembrane channel, initially within the bacterial bilayer. **b**, Alternative membrane fusion via a hemifusion intermediate with a diaphragm, preserving the transmembrane channel directionality.

Our results suggest a plausible solution to this conundrum via the formation of the hemi-fusion diaphragm (Fig. 5b). Here, the outer vesicle resembling an archaeon opens and widens –due to its chemical nature– allowing the inner vesicle to contact (fuse) highly unstable active edges along the rim of the external membrane, forming a hemifusion intermediate in the shape of a diaphragm. Finally, the bilayer isolating the two cavities, separates from the final structure generating an inner micelle. This mechanism successfully preserves the orientation of all transmembrane domains. Ultimately, the formation of an independent micelle at the core of the container vesicle requires a membrane fission event. In modern eukaryots, the ESCRT (Endosomal Sorting Complex Required for Transport) protein complex mediates fission events as well as plasma membrane resealing.^90,91^ This machinery was a contribution from the archaeal parental lineage.^92^ Accordingly, ESCRT-III filaments have been proposed to contribute to membrane remodeling during eukaryogenesis.^8,93^

The shared identity of bacterial and eukaryotic cell membranes has long posed a fundamental evolutionary question. Based on classical molecular dynamics simulations, we propose a plausible fusion mechanism between bacterial OMVs and the archaea plasma membrane —driven by acyclic bolalipid unstable edges— that could in principle explain the origin of the eukaryotic membrane composition (Fig. 6). This initial fusion event yields a composite cell membrane containing bacterial lipid patches, prone to expand via successive active-edge mediated fusions with newly produced OMVs.^94^ As bacterial OMVs became abundant in the cytoplasm of the host, they became targets of archaeal secretory protein systems giving rise to a primitive endomembrane system.^95^ Eventually, bacterial lipid synthesis overcomes its archaeal counterpart, owing to lower energetic cost^96^ and greater biophysical versatility (Fig. 1), allowing repeated cell divisions to progressively dilute the archaeal lipid signature. New cell generations would thus feature plasma membranes increasingly enriched in bacterial lipids and eventually lacking archaeal ones. At this stage, a primitive endomembrane system may have emerged, enabling the canonical membrane-fusion mechanisms that characterize eukaryotic cells. Our contribution from the biophysics perspective complements evolutionary research and brings light into an extraordinary fusion process that was driven by the chemical nature of the involved lipid species and their spatial interactions under confinement.

**Figure 6:**
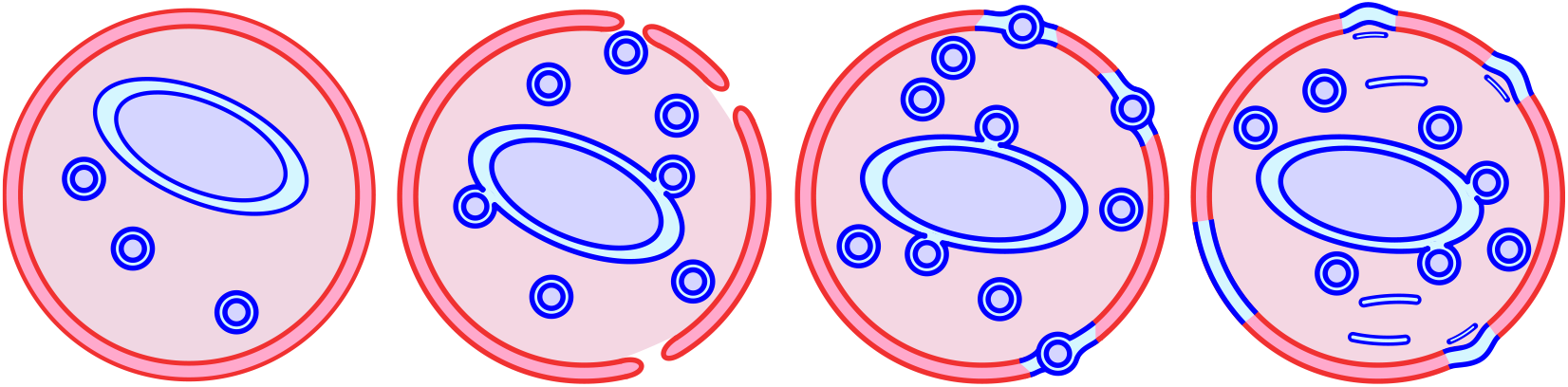
Initial steps toward eukaryotic membrane genesis via active-edge mediated fusion mechanism. Membranes from archeon and bacterium are respectively colored in red/pink and blue/cyan.

## Conclusions

The origin of the eukaryotic cell membrane represents a landmark transition in early life, yet it remains insufficiently explained. Traditional membrane fusion models have failed to explain full-fusion events between bacteria and archaea under the currently accepted hypothesis of eukaryogenesis. Using various high-performance computing techniques, we have directly observed archaeal-bacterial fusion via an unexpected, non-canonical mechanism that resolves previous inconsistencies, operates without traditional fusion pores, and requires no proteins to overcome free energy barriers. We show how confinement is the emergent property that drives archaeal and bacterial lipids to seamlessly merge into new stable structures, elegantly explaining the origin of the eukaryotic membrane. Our results reveal a fundamental incompatibility between the canonical fusion-pore model and the assumed bacterial-archaeal membrane ‘natural fusion’ mode. We observe a distinct pathway suggesting a counterintuitive fusion mechanism that unveils the emergence of the eukaryotic cell membrane. Overall, our work provides an interdisciplinary insight into a major evolutionary question, merging biophysics, computational modeling, and evolutionary biology.

## Author Contributions

LSM and MLM conceived the biological problem. DM performed the simulations and processed the trajectories. All authors analyzed the results and contributed to writing the manuscript.

## Conflicts of interest

There are no conflicts to declare.

## Data availability statement

Molecular dynamics files and examples related to this work are freely available through GitHub at https://github.com/diegomasone/bolalipids

## Acknowledgements

Supercomputing time for this work was provided by CCAD-UNC (Centro de Computación de Alto Desempeño de la Universidad Nacional de Córdoba). Grants from CONICET (PIP-0409CO) and ANPCyT (PICT2020-1897) are gratefully acknowledged as well as GPU hardware provided by the NVIDIA Corporation.

## Supporting Information Available

Supporting Information includes: contains snapshots of membrane fusion intermediate states; structures of DPPC, BOLA and BOLB vesicles under confinement; detailed steps along the edge-induced fusion mechanism; cholesterol inhibitory effects on edge-induced fusion; membrane density profiles; estimation of the hydration repulsion from single bolalipids membrane simulations; tables with statistics of important topological events; snapshots for DOPG:BOLB and DPPE:BOLB simulations.

## TOC Graphic

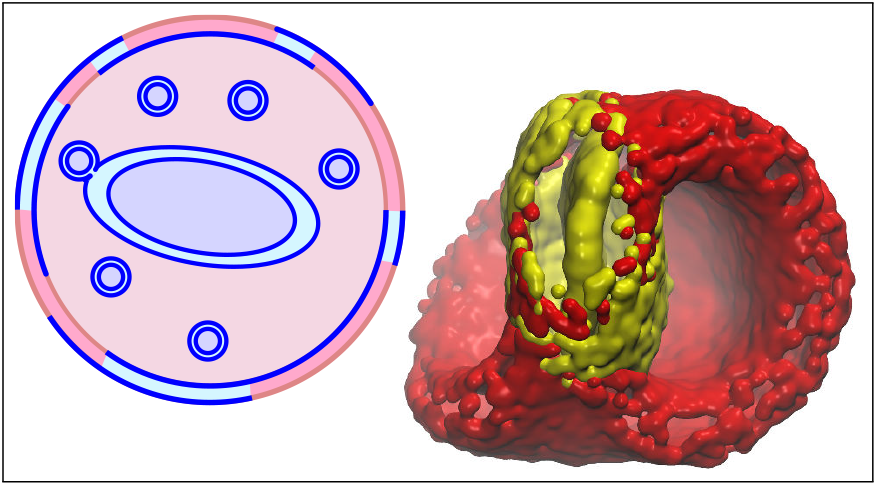

## Notes

https://github.com/diegomasone/bolalipids

### Competing Interest Statement

The authors have declared no competing interest.

## References

(1) Eme, L.; Spang, A.; Lombard, J.; Stairs, C. W.; Ettema, T. J. G. Archaea and the origin of eukaryotes. Nature Reviews Microbiology 2017, 15, 711–723.

(2) Gould, S. B. Membranes and evolution. Current Biology 2018, 28, R381–R385.

(3) Řezanka, T.; Kyselová, L.; Murphy, D. J. Archaeal lipids. Progress in Lipid Research 2023, 91, 101237.

(4) Chugunov, A. O.; Volynsky, P. E.; Krylov, N. A.; Boldyrev, I. A.; Efremov, R. G. Liquid but Durable: Molecular Dynamics Simulations Explain the Unique Properties of Archaeal-Like Membranes. Scientific Reports 2014, 4, 7462.

(5) Galimzyanov, T. R.; Kuzmin, P. I.; Pohl, P.; Akimov, S. A. Elastic deformations of bolalipid membranes. Soft Matter 2016, 12, 2357–2364.

(6) Sohlenkamp, C. Crossing the lipid divide. Journal of Biological Chemistry 2021, 297, 100859.

(7) Romero, H.; Aguilar, P. S.; Graña, M.; Langleib, M.; Gudiño, V.; Podbilewicz, B. Membrane fusion and fission during eukaryogenesis. Current Opinion in Cell Biology 2024, 86, 102321.

(8) Vosseberg, J.; van Hooff, J. J. E.; Köstlbacher, S.; Panagiotou, K.; Tamarit, D.; Ettema, T. J. G. The emerging view on the origin and early evolution of eukaryotic cells. Nature 2024, 633, 295–305.

(9) Gould, S. B.; Garg, S. G.; Martin, W. F. Bacterial Vesicle Secretion and the Evolutionary Origin of the Eukaryotic Endomembrane System. Trends in Microbiology 2016, 24, 525–534.

(10) Caforio, A.; Siliakus, M. F.; Exterkate, M.; Jain, S.; Jumde, V. R.; Andringa, R. L. H.; Kengen, S. W. M.; Minnaard, A. J.; Driessen, A. J. M.; van der Oost, J. Converting Escherichia coli into an archaebacterium with a hybrid heterochiral membrane. Proceedings of the National Academy of Sciences 2018, 115, 3704–3709.

(11) Devaux, P. F.; Morris, R. Transmembrane Asymmetry and Lateral Domains in Biological Membranes. Traffic 2004, 5, 241–246.

(12) Jahn, R.; Lang, T.; Südhof, T. C. Membrane Fusion. Cell 2003, 112, 519–533.

(13) Söllner, T. H. Intracellular and viral membrane fusion: a uniting mechanism. Current Opinion in Cell Biology 2004, 16, 429–435.

(14) Mayorga, L. S.; Tomes, C. N.; Belmonte, S. A. Acrosomal exocytosis, a special type of regulated secretion. IUBMB Life 2007, 59, 286–292.

(15) Chernomordik, L. V.; Kozlov, M. M.; Melikyan, G. B.; Abidor, I. G.; Markin, V. S.; Chizmadzhev, Y. A. The shape of lipid molecules and monolayer membrane fusion. Biochimica et Biophysica Acta (BBA) - Biomembranes 1985, 812, 643–655.

(16) Giraudo, C. G.; Hu, C.; You, D.; Slovic, A. M.; Mosharov, E. V.; Sulzer, D.; Melia, T. J.; Rothman, J. E. SNAREs can promote complete fusion and hemifusion as alternative outcomes. J Cell Biol 2005, 170, 249–260.

(17) Poojari, C. S.; Scherer, K. C.; Hub, J. S. Free energies of membrane stalk formation from a lipidomics perspective. Nature Communications 2021, 12, 6594.

(18) Van Der Spoel, D.; Lindahl, E.; Hess, B.; Groenhof, G.; Mark, A. E.; Berendsen, H. J. C. GROMACS: Fast, flexible, and free. Journal of Computational Chemistry 2005, 26, 1701–1718.

(19) Tribello, G.; Bonomi, M.; Branduardi, D.; Camilloni, C.; Bussi, G. PLUMED 2: New feathers for an old bird. Computer Physics Communications 2014, 185, 604–613.

(20) Marrink, S. J.; Risselada, H. J.; Yefimov, S.; Tieleman, D. P.; de Vries, A. H. The MARTINI Force Field: Coarse Grained Model for Biomolecular Simulations. J. Phys. Chem. B 2007, 111, 7812–7824.

(21) Jo, S.; Kim, T.; Iyer, V. G.; Im, W. CHARMM-GUI: A web-based graphical user interface for CHARMM. Journal of Computational Chemistry 2008, 29, 1859–1865.

(22) Wassenaar, T. A.; Ingolfsson, H. I.; Bockmann, R. A.; Tieleman, D. P.; Marrink, S. J. Computational Lipidomics with insane: A Versatile Tool for Generating Custom Membranes for Molecular Simulations. Journal of Chemical Theory and Computation 2015, 11, 2144–2155.

(23) Qi, Y.; Ingólfsson, H. I.; Cheng, X.; Lee, J.; Marrink, S. J.; Im, W. CHARMM-GUI Martini Maker for Coarse-Grained Simulations with the Martini Force Field. J. Chem. Theory Comput. 2015, 11, 4486–4494.

(24) Bussi, G.; Donadio, D.; Parrinello, M. Canonical sampling through velocity rescaling. The Journal of Chemical Physics 2007, 126, 014101.

(25) Seelig, A.; Seelig, J. Dynamic structure of fatty acyl chains in a phospholipid bilayer measured by deuterium magnetic resonance. Biochemistry 1974, 13, 4839–4845, PMID: 4371820.

(26) Parrinello, M.; Rahman, A. Polymorphic transitions in single crystals: A new molecular dynamics method. Journal of Applied Physics 1981, 52, 7182–7190.

(27) Smirnova, Y. G.; Risselada, H. J.; Müller, M. Thermodynamically reversible paths of the first fusion intermediate reveal an important role for membrane anchors of fusion proteins. Proceedings of the National Academy of Sciences 2019, 116, 2571–2576.

(28) Hub, J. S.; Awasthi, N. Probing a Continuous Polar Defect: A Reaction Coordinate for Pore Formation in Lipid Membranes. J. Chem. Theory Comput. 2017, 13, 2352–2366.

(29) Neale, C.; Pomes, R. Sampling errors in free energy simulations of small molecules in lipid bilayers. Biosimulations of lipid membranes coupled to experiments 2016, 1858, 2539–2548.

(30) Di Bartolo, A. L.; Masone, D. Synaptotagmin-1 C2B domains cooperatively stabilize the fusion stalk via a master-servant mechanism. Chem. Sci. 2022, 13, 3437–3446.

(31) Di Bartolo, A. L.; Tomes, C. N.; Mayorga, L. S.; Masone, D. Enhanced Expansion and Reduced Kiss-and-Run Events in Fusion Pores Steered by Synaptotagmin-1 C2B Domains. J. Chem. Theory Comput. 2022, 18, 4544–4554.

(32) Pearlman, D. A.; Kollman, P. A. The lag between the Hamiltonian and the system configuration in free energy perturbation calculations. J. Chem. Phys. 1989, 91, 7831– 7839.

(33) Brosio, G.; Rossi, G.; Bochicchio, D. Nanoparticle-induced biomembrane fusion: unraveling the effect of core size on stalk formation. Nanoscale Adv. 2023, 5, 4675–4680.

(34) Richardson, J. D.; Van Lehn, R. C. Free Energy Analysis of Peptide-Induced Pore Formation in Lipid Membranes by Bridging Atomistic and Coarse-Grained Simulations. J. Phys. Chem. B 2024, 128, 8737–8752.

(35) Awasthi, N.; Hub, J. S. Simulations of Pore Formation in Lipid Membranes: Reaction Coordinates, Convergence, Hysteresis, and Finite-Size Effects. J. Chem. Theory Comput. 2016, 12, 3261–3269.

(36) Roux, B. The calculation of the potential of mean force using computer simulations. Comput. Phys. Commun. 1995, 91, 275–282.

(37) Grossfield, A. WHAM: the weighted histogram analysis method. http://membrane.urmc.rochester.edu/wordpress/?page_id=126, version 2.0.9.1.

(38) Mayorga, L. S.; Masone, D. The Secret Ballet Inside Multivesicular Bodies. ACS Nano 2024, 18, 15651–15660.

(39) Boyd, K.; May, E. R. BUMPy: A Model-Independent Tool for Constructing Lipid Bilayers of Varying Curvature and Composition. J. Chem. Theory Comput. 2018, 14, 6642–6652.

(40) Mayorga, L. S.; Mascotti, M. L.; Bruininks, B. M. H.; Masone, D. Confinement Induces Morphological and Topological Transitions in Multivesicles. ACS Nano 2025, 19, 4515– 4527.

(41) Callan-Jones, A.; Sorre, B.; Bassereau, P. Curvature-driven lipid sorting in biomembranes. Cold Spring Harbor perspectives in biology 2011, 3, a004648.

(42) Risselada, H. J.; Marrink, S. J.; Muller, M. Curvature-Dependent Elastic Properties of Liquid-Ordered Domains Result in Inverted Domain Sorting on Uniaxially Compressed Vesicles. Phys. Rev. Lett. 2011, 106, 148102.

(43) Baoukina, S.; Ingólfsson, H. I.; Marrink, S. J.; Tieleman, D. P. Curvature-Induced Sorting of Lipids in Plasma Membrane Tethers. Adv. Theory Simul. 2018, 1, 1800034.

(44) Lipowsky, R. Coupling of bending and stretching deformations in vesicle membranes. Special issue in honour of Wolfgang Helfrich 2014, 208, 14–24.

(45) Turner, L.; Bitto, N. J.; Steer, D. L.; Lo, C.; D’Costa, K.; Ramm, G.; Shambrook, M.; Hill, A. F.; Ferrero, R. L.; Kaparakis-Liaskos, M. Helicobacter pylori Outer Membrane Vesicle Size Determines Their Mechanisms of Host Cell Entry and Protein Content. Frontiers in Immunology 2018, Volume 9 - 2018 .

(46) McCaig William, D.; Antonius, K.; Thanassi David, G. Production of Outer Membrane Vesicles and Outer Membrane Tubes by Francisella novicida. Journal of Bacteriology 2013, 195, 1120–1132.

(47) Furuyama, N.; Sircili, M. P. Outer Membrane Vesicles (OMVs) Produced by Gram-Negative Bacteria: Structure, Functions, Biogenesis, and Vaccine Application. BioMed Research Intl 2021, 2021, 1490732.

(48) Bruininks, B. M. H.; Thie, A. S.; Souza, P. C. T.; Wassenaar, T. A.; Faraji, S.; Marrink, S. J. Sequential Voxel-Based Leaflet Segmentation of Complex Lipid Morphologies. J. Chem. Theory Comput. 2021, 17, 7873–7885.

(49) Bruininks, B. M. H.; Wassenaar, T. A.; Vattulainen, I. Unbreaking Assemblies in Molecular Simulations with Periodic Boundaries. J. Chem. Inf. Model. 2023, 63, 3448–3452.

(50) Bruininks, B. M. H.; Vattulainen, I. Classification of containment hierarchy for point clouds in periodic space. bioRxiv 2025, 2025.08.06.668936.

(51) Humphrey, W.; Dalke, A.; Schulten, K. VMD - Visual Molecular Dynamics. Journal of Molecular Graphics 1996, 14, 33–38.

(52) Grace Development Team GRACE: GRaphing, Advanced Computation and Exploration of data. https://plasma-gate.weizmann.ac.il/Grace/, Last accessed: May 2024.

(53) Inkscape Project Inkscape. https://inkscape.org, Last accessed: May 2024.

(54) John W. Eaton S.H., David Bateman; Wehbring R. GNU Octave version 4.0.0 manual: a high-level interactive language for numerical computations; 2015.

(55) Stukowski, A. Visualization and analysis of atomistic simulation data with OVITO-the Open Visualization Tool. Modelling and Simulation in Materials Science and Engineering 2009, 18, 015012.

(56) Bulacu, M.; Périole, X.; Marrink, S. J. In Silico Design of Robust Bolalipid Membranes. Biomacromolecules 2012, 13, 196–205.

(57) Smirnova, Y. G.; Aeffner, S.; Risselada, H. J.; Salditt, T.; Marrink, S. J.; Muller, M.; Knecht, V. Interbilayer repulsion forces between tension-free lipid bilayers from simulation. Soft Matter 2013, 9, 10705–10718.

(58) Parsegian, V. A.; Fuller, N.; Rand, R. P. Measured work of deformation and repulsion of lecithin bilayers. Proceedings of the National Academy of Sciences 1979, 76, 2750–2754.

(59) Smirnova, Y. G.; Marrink, S.-J.; Lipowsky, R.; Knecht, V. Solvent-exposed tails as prestalk transition states for membrane fusion at low hydration. Journal of the American Chemical Society 2010, 132, 6710–6718.

(60) Sharma, S.; Lindau, M. The fusion pore, 60 years after the first cartoon. FEBS Lett 2018, 592, 3542–3562.

(61) Sagan, L. On the origin of mitosing cells. Journal of Theoretical Biology 1967, 14, 225–IN6.

(62) Knapp, B.; Ospina, L.; Deane, C. M. Avoiding False Positive Conclusions in Molecular Simulation: The Importance of Replicas. J. Chem. Theory Comput. 2018, 14, 6127– 6138.

(63) Kozlovsky, Y.; Chernomordik, L. V.; Kozlov, M. M. Lipid Intermediates in Membrane Fusion: Formation, Structure, and Decay of Hemifusion Diaphragm. Biophysical Journal 2002, 83, 2634–2651.

(64) Knorr, R. L.; Mizushima, N.; Dimova, R. Fusion and scission of membranes: Ubiquitous topological transformations in cells. Traffic 2017, 18, 758–761.

(65) Weirich, K. L.; Israelachvili, J. N.; Fygenson, D. K. Bilayer Edges Catalyze Supported Lipid Bilayer Formation. Biophysical Journal 2010, 98, 85–92.

(66) Chlanda, P.; Mekhedov, E.; Waters, H.; Schwartz, C. L.; Fischer, E. R.; Ryham, R. J.; Cohen, F. S.; Blank, P. S.; Zimmerberg, J. The hemifusion structure induced by influenza virus haemagglutinin is determined by physical properties of the target membranes. Nature microbiology 2016, 1, 16050.

(67) Risselada, H. J.; Marelli, G.; Fuhrmans, M.; Smirnova, Y. G.; Grubmuller, H.; Marrink, S. J.; Muller, M. Line-Tension Controlled Mechanism for Influenza Fusion. PLOS ONE 2012, 7, 1–14.

(68) Bartolo, D.; Lautaro, A.; Caparotta, M.; Polo, L. M.; Masone, D. Myomerger Induces Membrane Hemifusion and Regulates Fusion Pore Expansion. Biochemistry 2024, 63, 815–826.

(69) Katsov, K.; Müller, M.; Schick, M. Field Theoretic Study of Bilayer Membrane Fusion. Hemifusion Mechanism. Biophysical Journal 2004, 87, 3277–3290.

(70) Müller, M.; Katsov, K.; Schick, M. A New Mechanism of Model Membrane Fusion Determined from Monte Carlo Simulation. Biophysical Journal 2003, 85, 1611–1623.

(71) Katsov, K.; Müller, M.; Schick, M. Field Theoretic Study of Bilayer Membrane Fusion: II. Mechanism of a Stalk-Hole Complex. Biophysical Journal 2006, 90, 915–926.

(72) Zhen, Y.; Radulovic, M.; Vietri, M.; Stenmark, H. Sealing holes in cellular membranes. The EMBO Journal 2021, 40, e106922.

(73) Hämälistö, S.; Stahl, J. L.; Favaro, E.; Yang, Q.; Liu, B.; Christoffersen, L.; Loos, B.; Guasch Boldú, C.; Joyce, J. A.; Reinheckel, T. et al. Spatially and temporally defined lysosomal leakage facilitates mitotic chromosome segregation. Nature Communications 2020, 11, 229.

(74) Shangguan, T.; Alford, D.; Bentz, J. Influenza Virus-Liposome Lipid Mixing Is Leaky and Largely Insensitive to the Material Properties of the Target Membrane. Biochemistry 1996, 35, 4956–4965.

(75) Risselada, H.; Smirnova, Y.; Grubmüller, H. Free Energy Landscape of Rim-Pore Expansion in Membrane Fusion. Biophysical Journal 2014, 107, 2287–2295.

(76) Spencer, R. K. W.; Smirnova, Y. G.; Soleimani, A.; Müller, M. Transient pores in hemifusion diaphragms. Biophysical Journal 2024, 123, 2455–2475.

(77) Golani, G.; Schwarz, U. S. High curvature promotes fusion of lipid membranes: Predictions from continuum elastic theory. Biophysical Journal 2023, 122, 1868–1882.

(78) Nikolaus, J.; Stöckl, M.; Langosch, D.; Volkmer, R.; Herrmann, A. Direct Visualization of Large and Protein-Free Hemifusion Diaphragms. Biophysical Journal 2010, 98, 1192–1199.

(79) Mattie, S.; McNally, E. K.; Karim, M. A.; Vali, H.; Brett, C. L. How and why intralumenal membrane fragments form during vacuolar lysosome fusion. MBoC 2017, 28, 309–321.

(80) Chanturiya, A.; Scaria, P.; Woodle, M. C. The Role of Membrane Lateral Tension in Calcium-Induced Membrane Fusion. The Journal of Membrane Biology 2000, 176, 67–75.

(81) van Tilburg; Marco P.A.; Marrink, S. J.; König, M.; Grünewald, F. Shocker - A Molecular Dynamics Protocol and Tool for Accelerating and Analyzing the Effects of Osmotic Shocks. J. Chem. Theory Comput. 2024, 20, 212–223.

(82) Bruckner, R. J.; Mansy, S. S.; Ricardo, A.; Mahadevan, L.; Szostak, J. W. Flip-Flop-Induced Relaxation of Bending Energy: Implications for Membrane Remodeling. Biophysical Journal 2009, 97, 3113–3122.

(83) Sreekumari, A.; Lipowsky, R. Large stress asymmetries of lipid bilayers and nanovesicles generate lipid flip-flops and bilayer instabilities. Soft Matter 2022, 18, 6066–6078.

(84) Franke, J. D. Developmental Biology in Prokaryotes and Lower Eukaryotes; Springer International Publishing: Cham, 2021; pp 215–227.

(85) Mangiarotti, A.; Sabri, E.; Schmidt, K. V.; Hoffmann, C.; Milovanovic, D.; Lipowsky, R.; Dimova, R. Lipid packing and cholesterol content regulate membrane wetting and remodeling by biomolecular condensates. Nature Communications 2025, 16, 2756.

(86) Fábián, B.; Vattulainen, I.; Javanainen, M. Protein Crowding and Cholesterol Increase Cell Membrane Viscosity in a Temperature Dependent Manner. J. Chem. Theory Comput. 2023, 19, 2630–2643.

(87) Kreutzberger, A. B.; Kiessling, V.; Tamm, L. High Cholesterol Obviates a Prolonged Hemifusion Intermediate in Fast SNARE-Mediated Membrane Fusion. Biophysical Journal 2015, 109, 319–329.

(88) Huang, Z.; London, E. Cholesterol lipids and cholesterol-containing lipid rafts in bacteria. Properties and Functions of Cholesterol 2016, 199, 11–16.

(89) Brown, A. J.; Galea, A. M. Cholesterol as an evolutionary response to living with oxygen. Evol 2010, 64, 2179–2183.

(90) Harker-Kirschneck, L.; Hafner, A. E.; Yao, T.; Vanhille-Campos, C.; Jiang, X.; Pulschen, A.; Hurtig, F.; Hryniuk, D.; Culley, S.; Henriques, R. et al. Physical mechanisms of ESCRT-III-driven cell division. Proceedings of the National Academy of Sciences 2022, 119, e2107763119.

(91) Pfitzner, A.-K.; Mercier, V.; Jiang, X.; von Filseck, J. M.; Baum, B.; Šarić, A.; Roux, A. An ESCRT-III polymerization sequence drives membrane deformation and fission. Cell 2020, 182, 1140–1155.

(92) Lindås, A.-C.; Karlsson, E. A.; Lindgren, M. T.; Ettema, T. J. G.; Bernander, R. A unique cell division machinery in the Archaea. Proceedings of the National Academy of Sciences 2008, 105, 18942–18946.

(93) Melnikov, N.; Junglas, B.; Halbi, G.; Nachmias, D.; Zerbib, E.; Gueta, N.; Upcher, A.; Zalk, R.; Sachse, C.; Bernheim-Groswasser, A. The Asgard archaeal ESCRT-III system forms helical filaments and remodels eukaryotic-like membranes. The EMBO journal 2025, 44, 665–681.

(94) Deatherage Brooke, L.; Cookson Brad, T. Membrane Vesicle Release in Bacteria, Eukaryotes, and Archaea: a Conserved yet Underappreciated Aspect of Microbial Life. Infection and Immunity 2012, 80, 1948–1957.

(95) Ellen, A. F.; Zolghadr, B.; Driessen, A. M. J.; Albers, S.-V. Shaping the Archaeal Cell Envelope. Archaea 2010, 2010, 608243.

(96) Mencía, M. The archaeal-bacterial lipid divide, could a distinct lateral proton route hold the answer? Biology Direct 2020, 15, 7.

