## supporting information for "Unconventional Fusion Mechanism at the Origin of Eukaryotic Membranes"

### Supporting Information: Unconventional Fusion Mechanism at the Origin of Eukaryotic Membranes

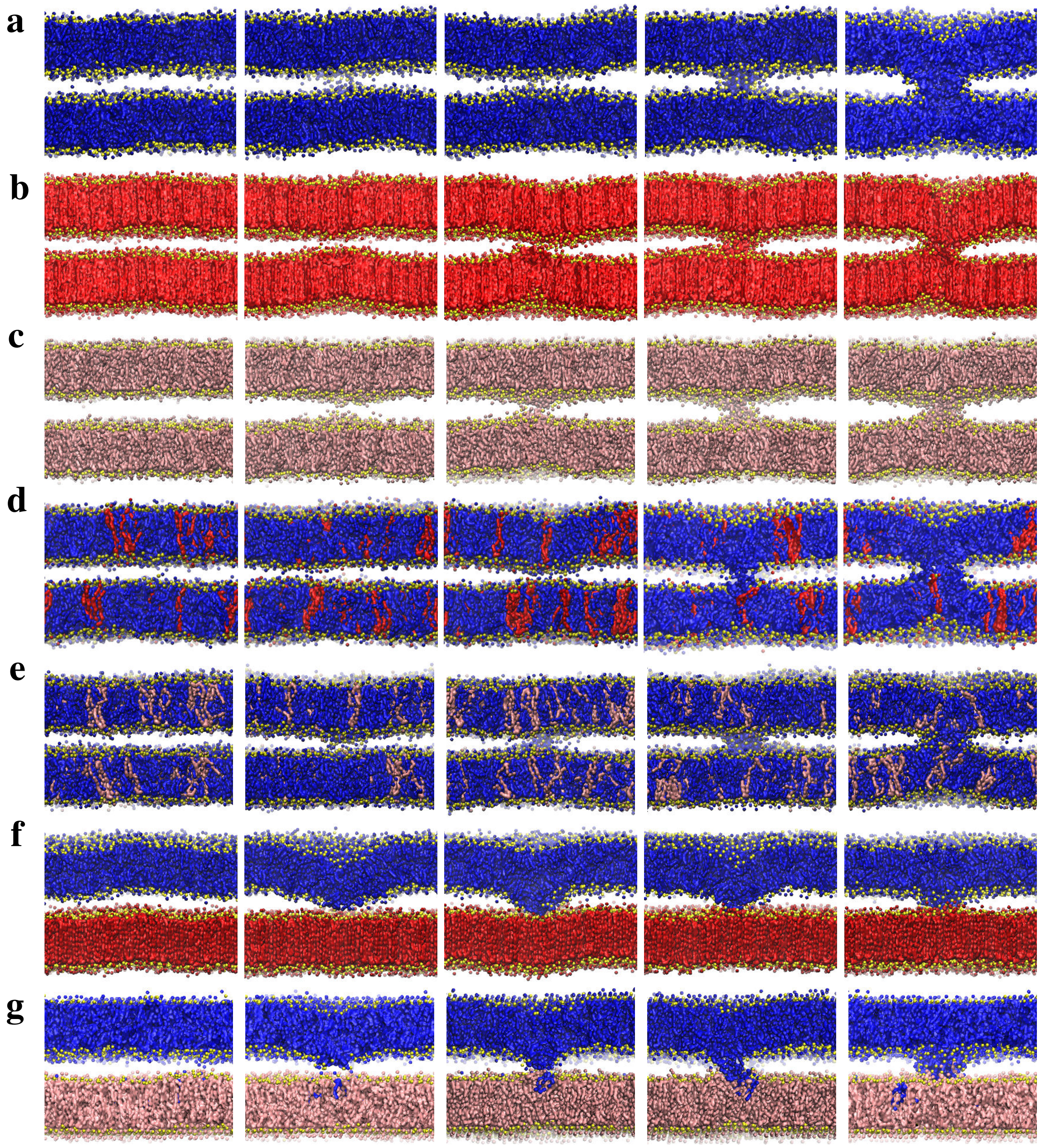


Figure S1: **Membrane fusion with intermediate transient states.** **a**, Pure DPPC. **b**, Pure BOLA. **c**, Pure BOLB. **d**, DPPC:BOLA (90:10). **e**, DPPC:BOLB (90:10). **f**, DPPC:BOLA (sep). **g**, DPPC:BOLB (sep). DPPC is blue, BOLA is red, BOLB is pink and phosphate groups (PO4, PO1 and PO2) are yellow. Water molecules are not shown.

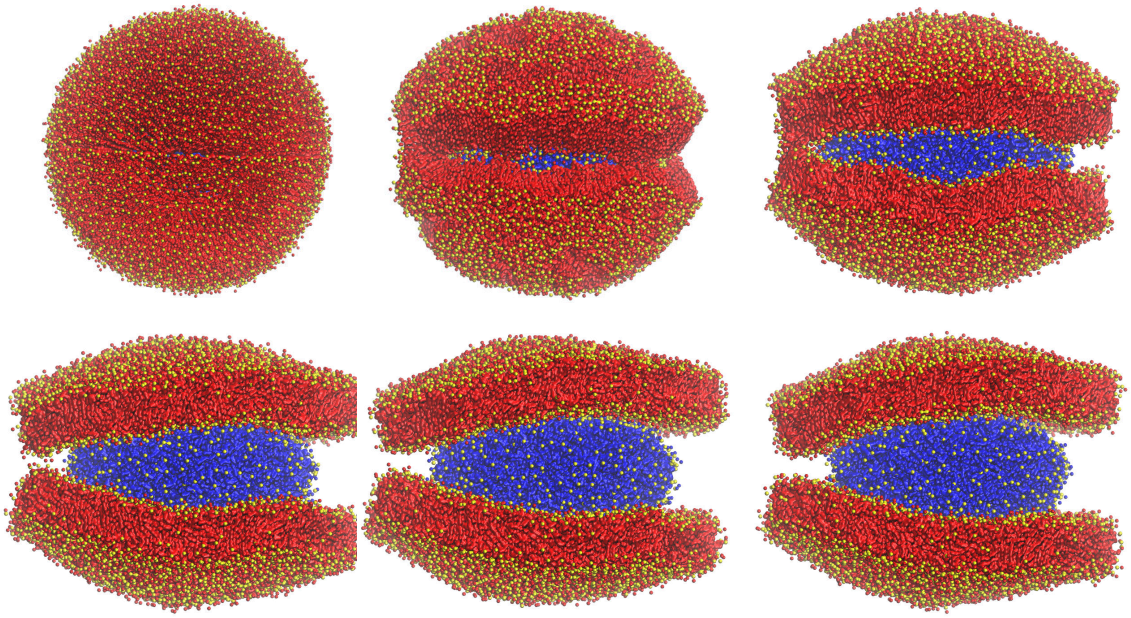

Figure S2: **DPPC(inner):BOLA(outer).** Temporal evolution over 10$\mu$s. DPPC is blue, BOLA is red. All phosphate groups are yellow. Water molecules are not shown.

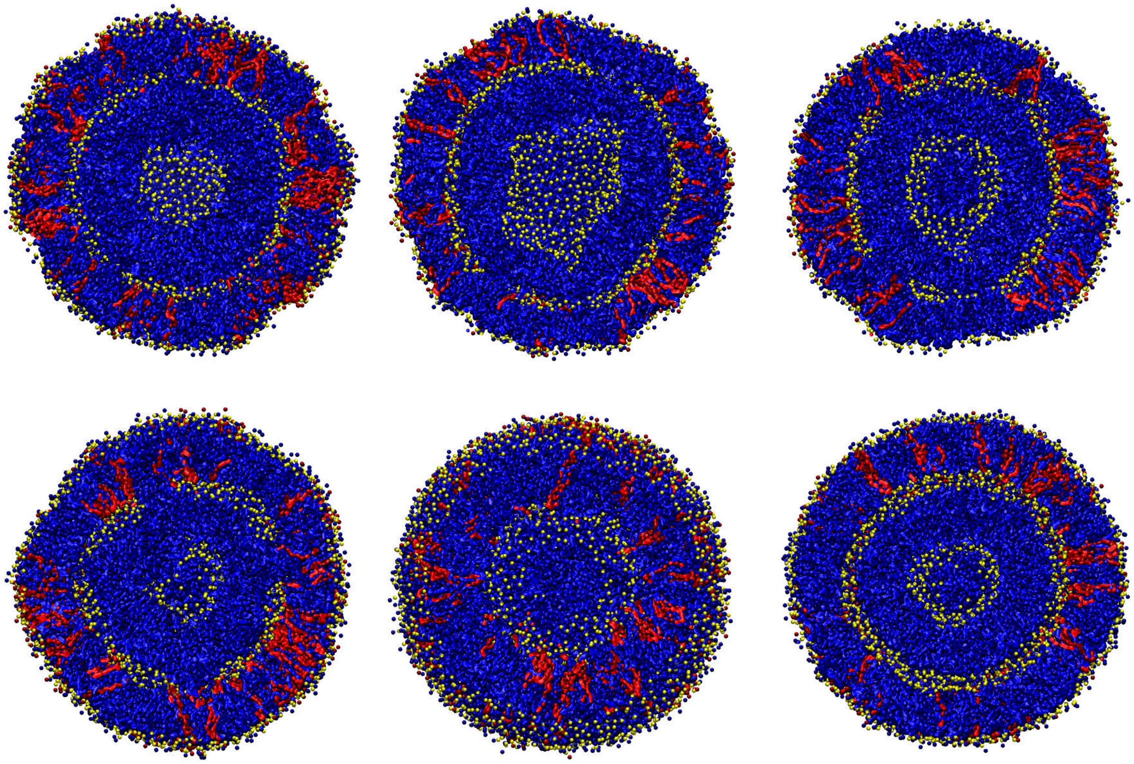

Figure S3: **DPPC:BOLA (90:10).** Temporal evolution over 10$\mu$s. DPPC is blue, BOLA is red. All phosphate groups are yellow. Water molecules are not shown.

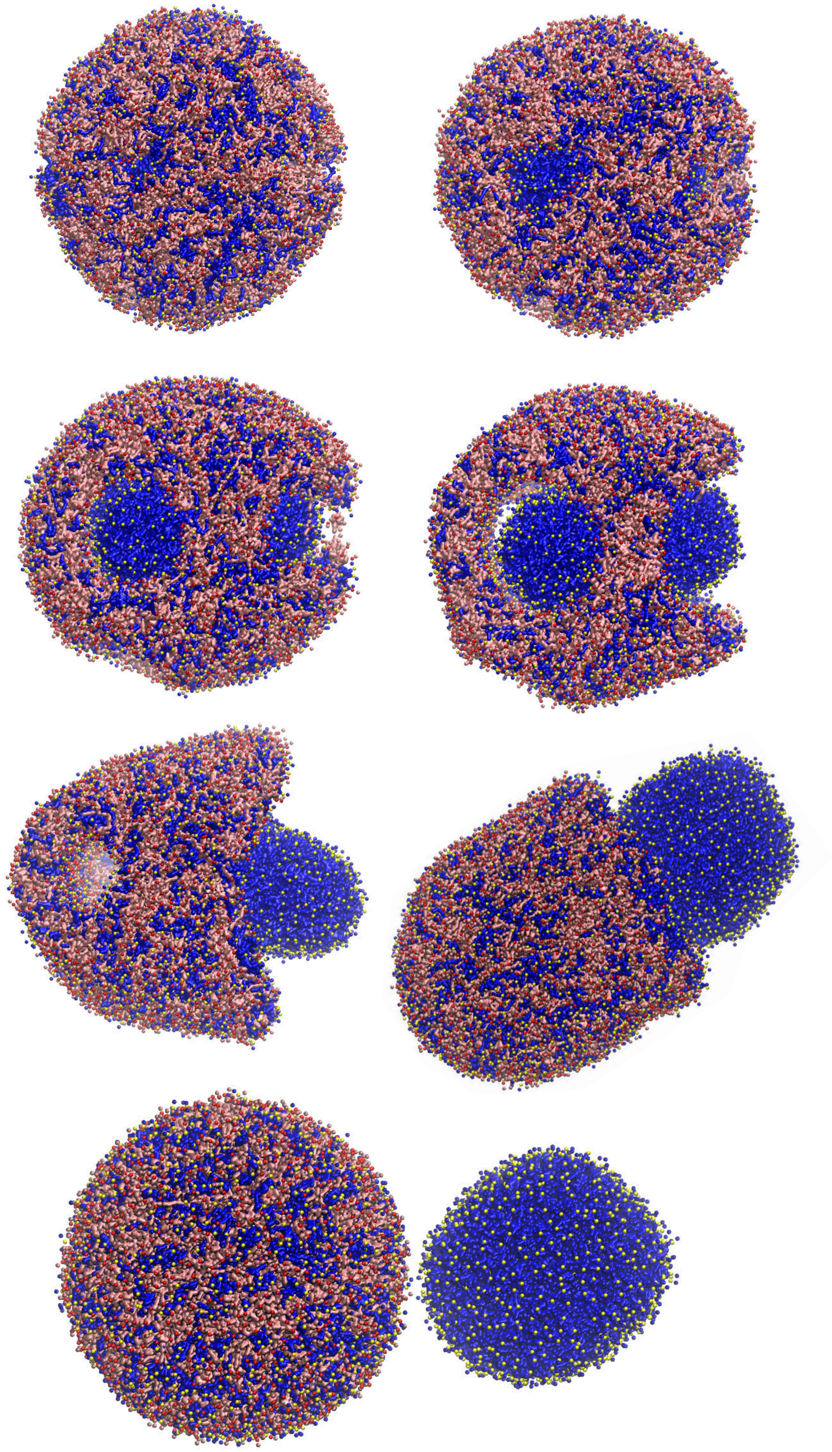

Figure S4: **DPPC:BOLB (50:50).** Temporal evolution over 10$\mu$s. DPPC is blue, BOLB is pink. PO4 phosphate groups are yellow and PO1,PO2 are yellow. Water molecules are not shown.

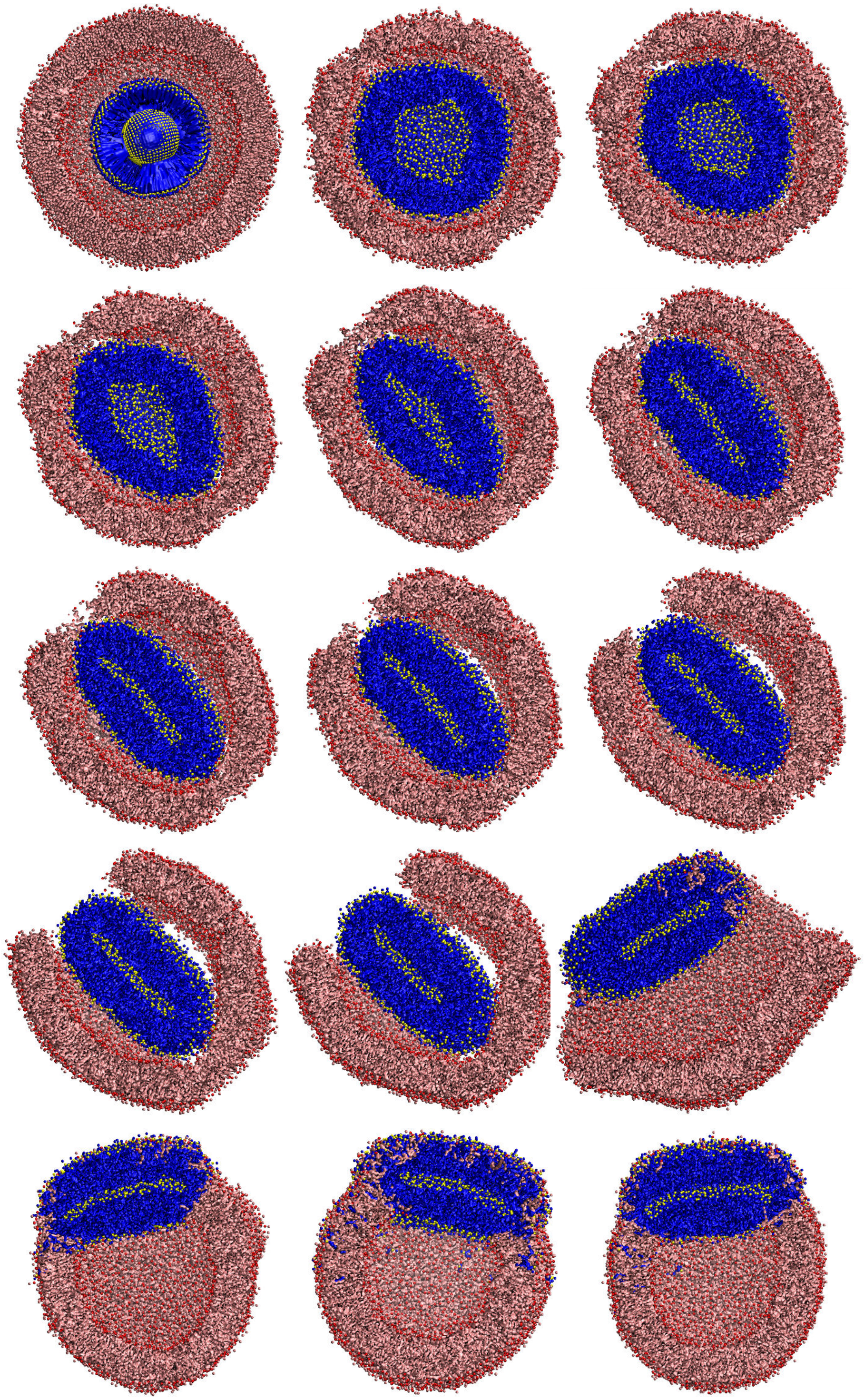

Figure S5: **DPPC(inner):BOLB(outer).** Hemifusion diaphragm formation via active-edge membrane fusion. Representative temporal evolution of 10 out of 10 trajectories in the $\mu$s-length scale. DPPC is blue, BOLB is pink. PO4 phosphate groups are yellow and PO1,PO2 are red. Water molecules are not shown.

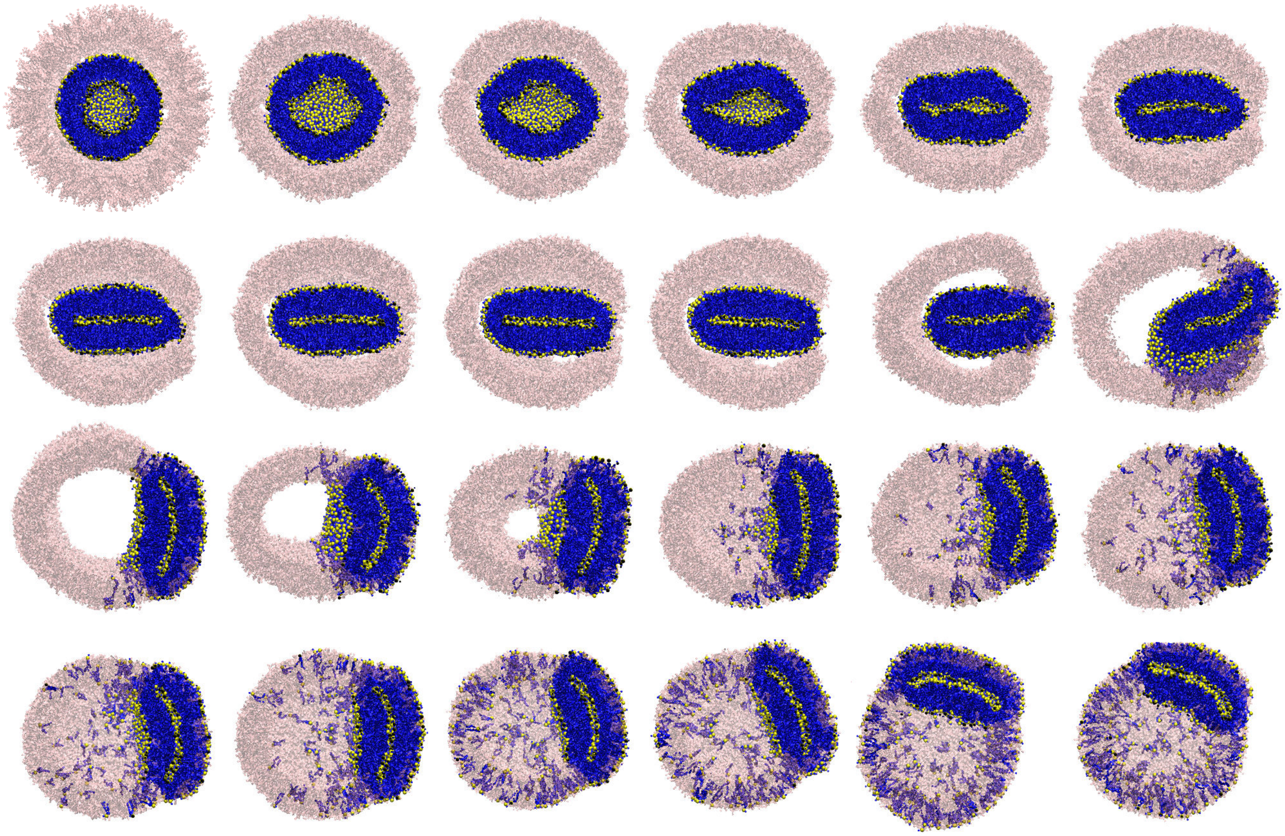

Figure S6: **Transient structures during equilibration and first steps of production runs.** DPPC is blue with PO4 beads in yellow. BOLB is transparent-pink.

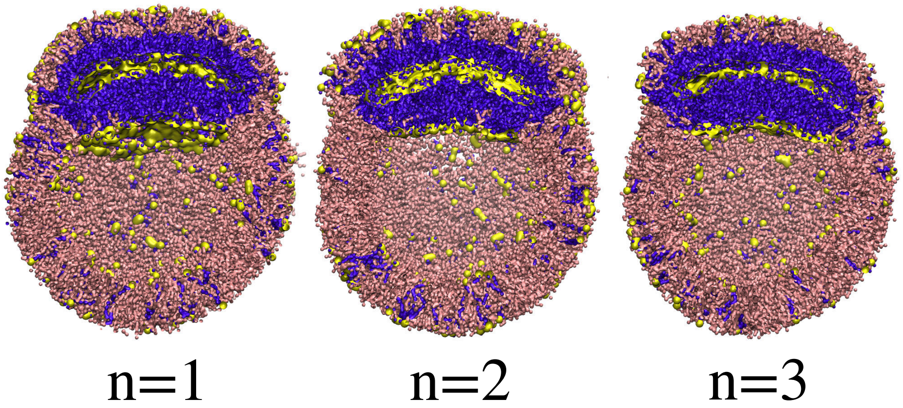


Figure S7: **DPPE:BOLB vesicle-in-vesicle.** Production-runs outcomes from 3 independent simulations showing the formation of a hemifusion intermediate. BOLB is pink, DPPE is violet with PO4 beads in yellow. Water molecules are not shown.

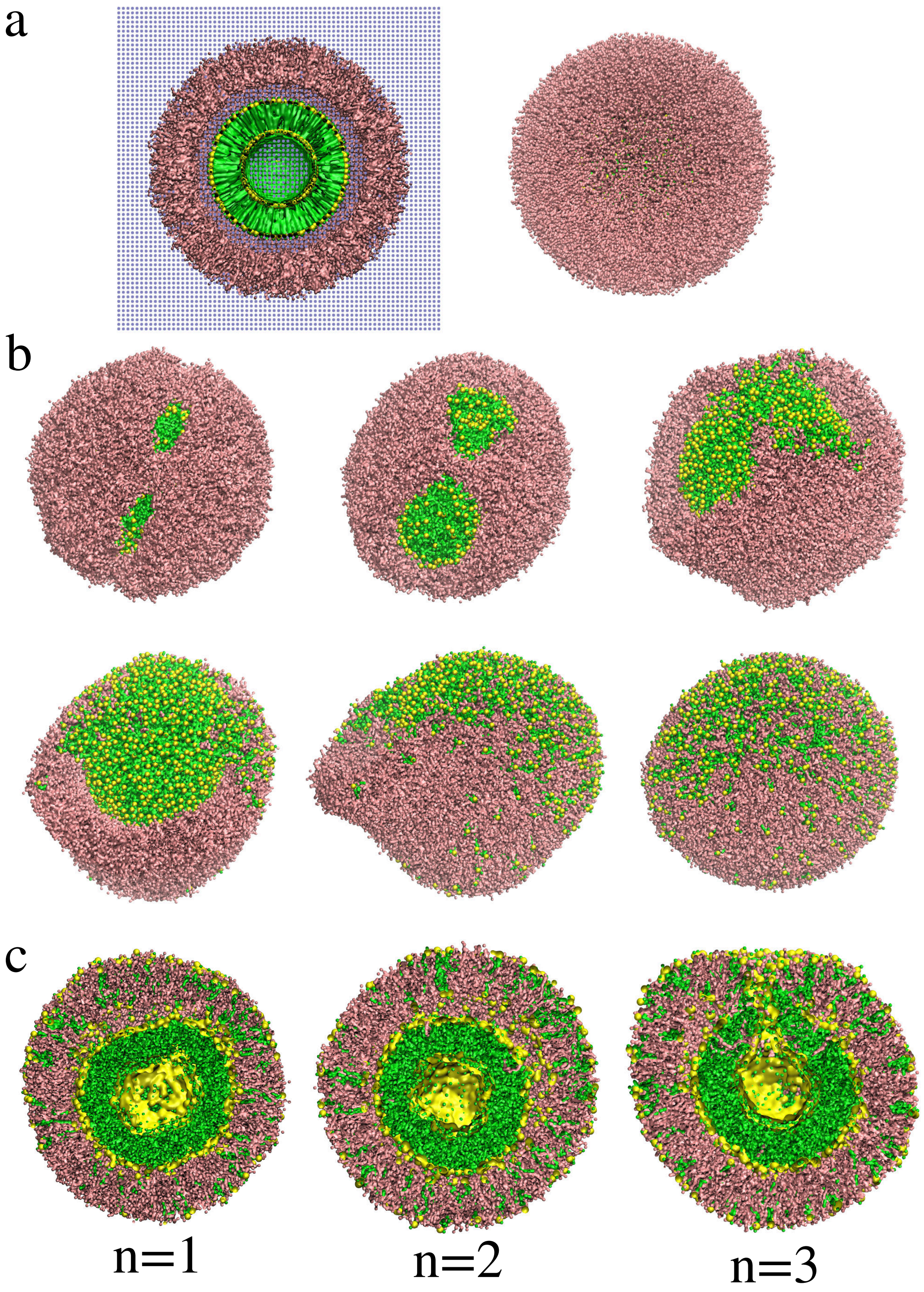


Figure S8: **DOPG:BOLB vesicle-in-vesicle. a,** Initial set-up. **b,** Transient states during equilibration. **c,** Production-runs outcomes from 3 independent simulations. BOLB is pink, DOPG is green with PO4 beads in yellow. Water molecules are only shown for the initial panel.

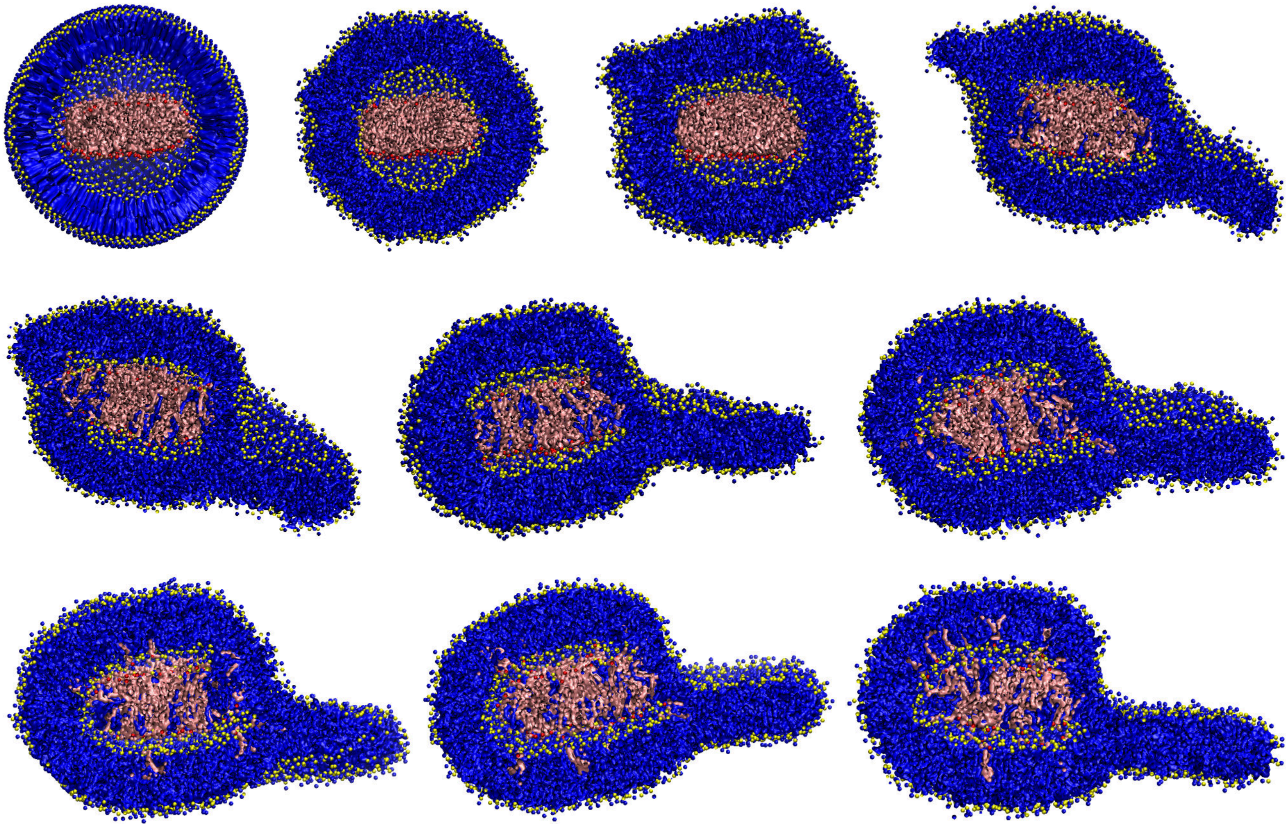

Figure S9: **BOLB edge-induced fusion.** Temporal sequence of a BOLB micelle confined by a DPPC vesicle (cross-view). Lipid excess form an external bleb. DPPC is blue, BOLB is pink. PO4 phosphate groups are yellow and PO1,PO2 are red.

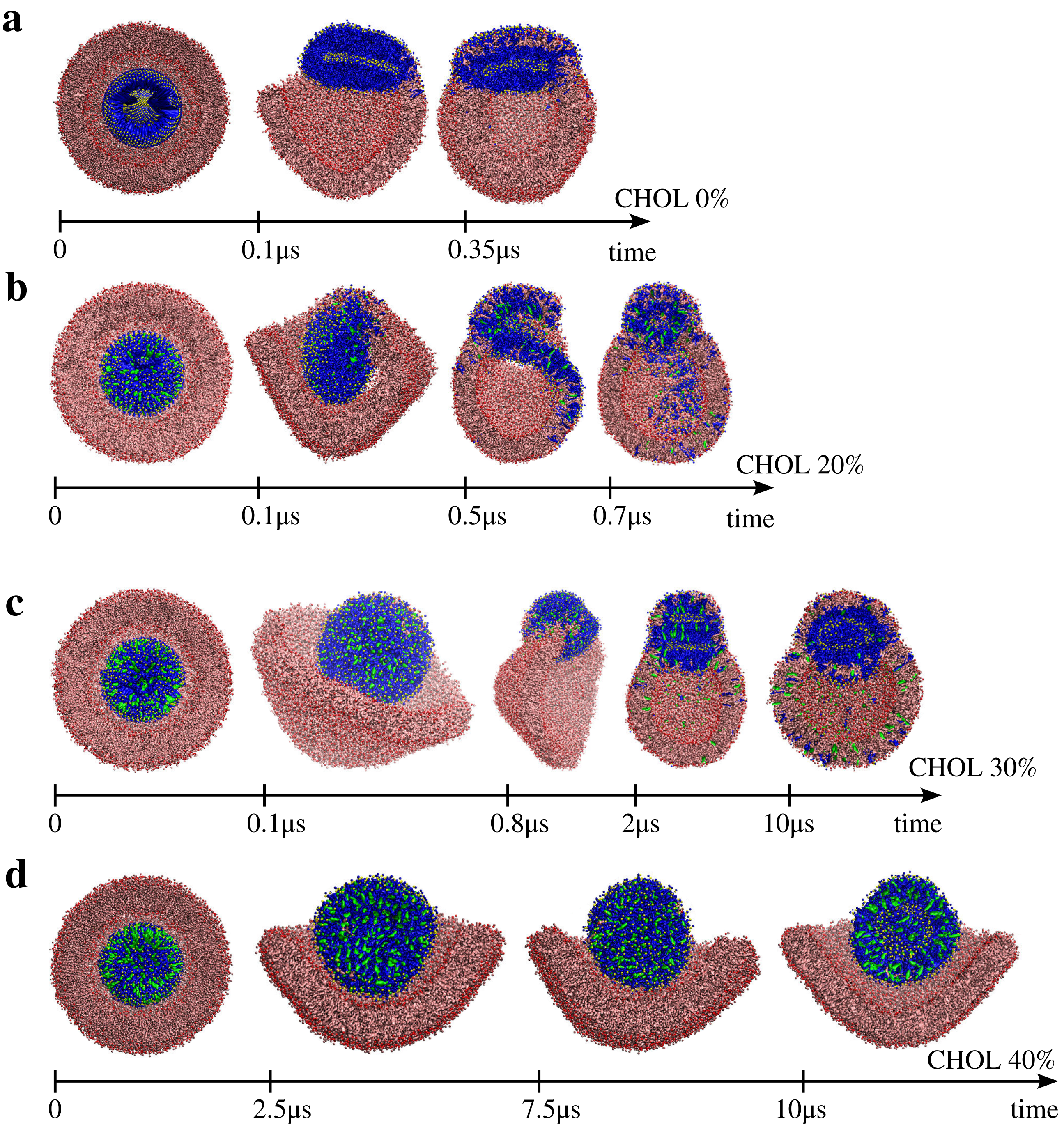


Figure S10: **Cholesterol inhibitory effects on active-edge mediated fusion until the first hemifusion diaphragm.** **a**, 0% cholesterol. **b**, 20% cholesterol. **c**, 30% cholesterol. **d**, 40% cholesterol. DPPC is blue, BOLB is pink and cholesterol is green. PO4 phosphate groups are yellow and PO1,PO2 are red. Water molecules are not shown.

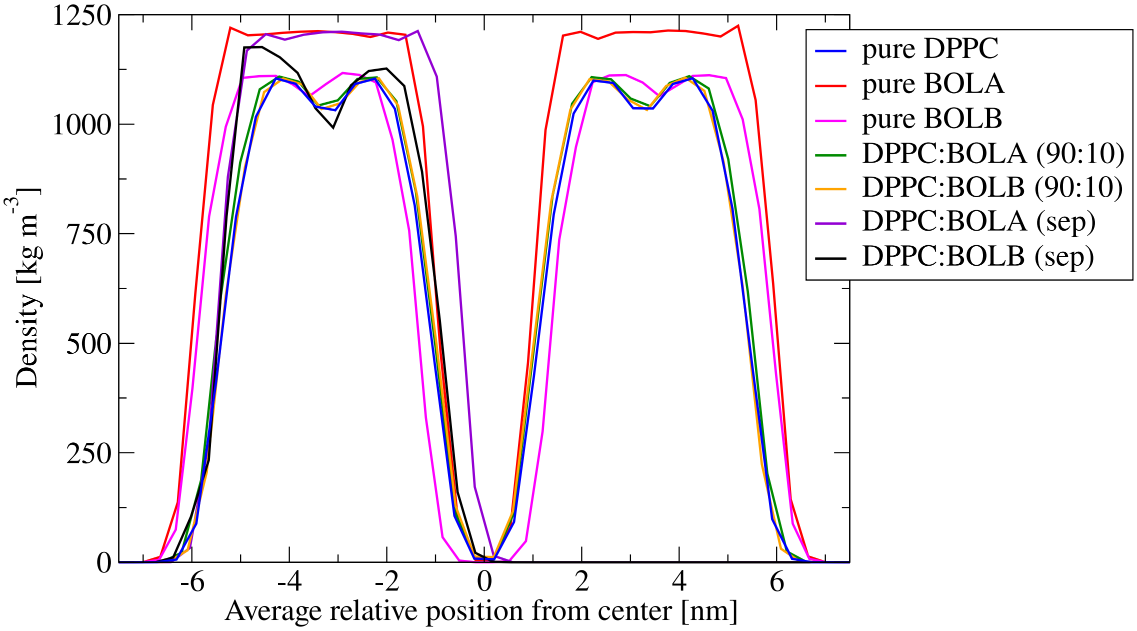


Figure S11: **Density profiles along the normal axis to the bilayers.** Calculated over all systems listed in table 1. These profiles show that the initially equilibrated inter-membrane distance is $\sim$ 4nm for all systems, before inducing the hemifusion stalk. Small variations are due to differences in lipid composition, producing small variations in the bilayer thickness.

#### Hydration repulsion from single-membrane simulations

To support conclusions from the stalk formation free energy profiles obtained via umbrella sampling, we calculated the hydration repulsion between a single bolalipid membrane and its periodic image. The procedure follows the framework developed by Smirnova *et al.*[1] in agreement with previous experimental measurements by Rawicz *et al.*[2], and allows for direct extraction of the disjoining pressure $P$ from unbiased molecular dynamics simulations.

First, a single 512 BOLB ∼16x18nm patch was hydrated with a number of water molecules that mimics the initial inter-membrane distance of $\sim 4nm$ in the double membrane systems used for stalk formation in biased simulations. This system was minimized, equilibrated and subsequently simulated in the $NPT$ ensemble with semi-isotropic pressure coupling at 1bar, so that the membrane remained tension-free at full hydration. Production runs were conducted for 1$\mu$s (in the Martini's coarse-grained space). The area per lipid $A$ was computed from the lateral box area divided by the amount of lipids in the membrane (monolayer) $N = 512$. The reference area per lipid $A_0$ was taken from the zero-tension, fully hydrated state.

Second, water thickness $d_w$ was estimated using the box dimension $L_z$ in the membrane normal direction and the headgroup-headgroup distance $d_{hh}$ of the membrane (PO1 and PO2 beads). The membrane thickness $d_{hh}$ was obtained from density profiles of the phosphate headgroups using `gmx density`. The two density maxima corresponding to the upper and lower surfaces were identified, and their separation $d_{hh}$ was measured, see figure S9. The inter-membrane water thickness then follows as: $d_w = L_z - d_{hh} = 8.85nm - 4.20nm = 4.65nm$.

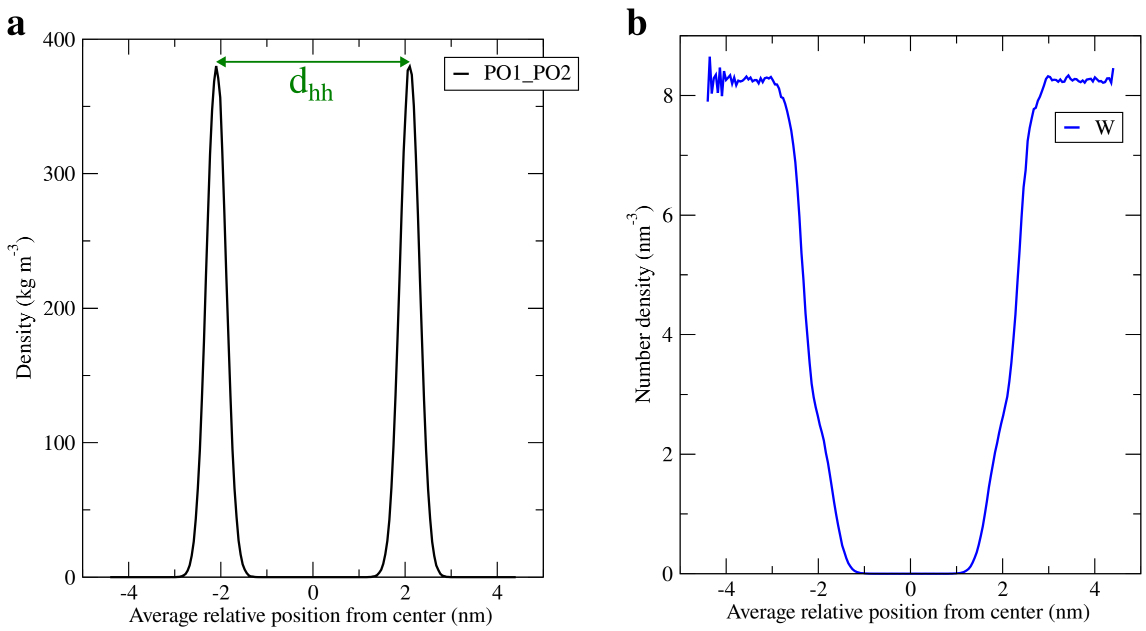


Figure S12: **Density profiles along the normal axis to the single bolalipid membrane. a,** Phosphate groups density. **b,** Water density.

Third, the bilayer area compressibility modulus $K_A$ was estimated to relate structural changes to the disjoining pressure. $K_A$ was determined from a set of semi-isotropic $NPT$ simulations performed under varying lateral reference pressures (stretching and compressing). The mean membrane tension $\Sigma$ was calculated from the pressure tensor according to:

$$\Sigma = \left\langle L_z \left( P_{zz} - \frac{P_{xx} + P_{yy}}{2} \right) \right\rangle$$

where $P_{ii}$ are the diagonal elements of the pressure tensor. The relationship between tension and area strain is:

$$\Sigma = K_A \frac{\Delta A}{A_0} \tag{1}$$

where $\Delta A = A - A_0$. Linear regression of $\Sigma$ against $\Delta A/A_0$ yields $K_A \sim 615.7 \pm 72.9 mN/m$, as shown in figure S10. The uncertainty in $K_A$ is the standard error of the slope from a least squares fitting.

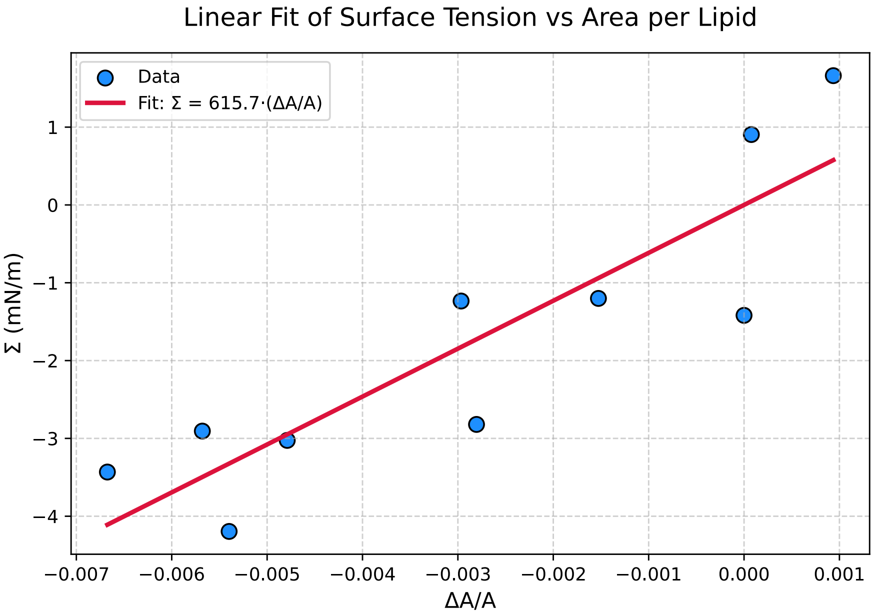


Figure S13: **Estimation of the area compressibility modulus $K_A$ for a single bolalipid membrane.**

Finally, equation 2 allows for the calculation of the hydration repulsion pressure directly from equilibrium simulations of a single bilayer with its periodic image, without requiring explicit force application and in terms of measurable simulation observables[1], yielding $logP \sim 5\text{mN/m}$, in agreement with reported values for other model membranes[1,3].

$$P = \frac{K_A}{d_w}\left(1 - \frac{A}{A_0}\right) \tag{2}$$

Table S1: Fast mechanism. Mean time to full fusion: MTFF=21.81$\mu$s

| Simulation run # | Hemifusion diaphragm forms [$\mu$s] | Rim pore opens in hemifusion diaphragm [$\mu$s] | Fission event separates inner micelle [$\mu$s] |
|---|---|---|---|
| 1 | 0.35 | 11.42 | 11.70 |
| 2 | 0.32 | 25.76 | 25.87 |
| 3 | 0.24 | 7.29 | 7.75 |
| 4 | 0.25 | 30.85 | 31.48 |
| 5 | 0.37 | 25.34 | 26.03 |
| 6 | 0.52 | 28.93 | 29.05 |
| 7 | 0.33 | 19.31 | 19.67 |
| 8 | 0.29 | 19.63 | 19.71 |
| 9 | 0.28 | 26.38 | 26.45 |
| 10 | 0.50 | 20.25 | 20.37 |

Table S2: Slow mechanism. Mean time to full fusion: MTFF=27.65$\mu$s

| Simulation run # | Rim pore opens in hemifusion diaphragm [$\mu$s] | Bleb-like micelle is completely reabsorbed [$\mu$s] |
|---|---|---|
| 11 | 19.94 | 21.26 |
| 12 | 29.32 | 31.37 |
| 13 | 41.47 | 44.26 |
| 14 | 21.78 | 24.41 |
| 15 | 20.05 | 21.99 |
| 16 | 36.43 | 38.36 |
| 17 | 21.36 | 23.99 |
| 18 | 26.29 | 27.54 |
| 19 | 19.60 | 22.04 |
| 20 | 19.80 | 21.30 |

### References


(1): Smirnova, Y. G.; Aeffner, S.; Risselada, H. J.; Salditt, T.; Marrink, S. J.; Muller, M.; Knecht, V. Interbilayer repulsion forces between tension-free lipid bilayers from simulation. *Soft Matter* **2013**, 9, 10705-10718.

(2): Rawicz, W.; Olbrich, K. C.; McIntosh, T.; Needham, D.; Evans, E. Effect of chain length and unsaturation on elasticity of lipid bilayers. *Biophysical journal* **2000**, 79(1), 328-339.

(3): Parsegian, V. A.; Fuller, N.; Rand, R. P. Measured work of deformation and repulsion of lecithin bilayers. *Proceedings of the National Academy of Sciences* **1979**, 76, 2750–2754.
